# A room-temperature ⁸⁹Zr⁴⁺ radiolabelling strategy for small extracellular vesicles with enhanced plasma stability for PET Imaging

**DOI:** 10.64898/2026.01.21.700868

**Authors:** Arnab Banerjee, Ivanna Hrynchak, Carlos Jesus, José Sereno, Tânia Martins-Marques, Magda Silva, Maria João Ferreira, Henrique Girao, Antero Abrunhosa, Lino Ferreira

## Abstract

Both for diagnostic purposes and regenerative medicine, it is essential to develop advanced imaging platforms capable of tracking the biodistribution of small extracellular vesicles (sEVs), as current methods are limited by inadequate resolution and sensitivity. In this study, we introduce a novel labeling strategy utilizing the radioisotope zirconium-89 (^89^Zr), which boasts a half-life of 78.4 h and is cost-effective to produce. To achieve this, we designed a new chelator tailored for ^89^Zr^4+^ that offers enhanced stability compared to the conventional deferoxamine (DFO). This chelator forms a robust complex with ^89^Zr^4+^ at room temperature, suitable for sEV labeling for PET imaging applications. The radiolabeling process involved a two-step procedure: first, conjugation of the chelator to the sEVs, and second, radiolabeling with ^89^Zr^4+^. The resulting sEV-L1-Zr demonstrated a radiochemical yield of approximately 60% and maintained around 80% stability in plasma over seven days. Importantly, our modifications did not alter the morphology, surface protein composition, internal RNA content, or bioactivity of the sEVs. We successfully visualized sEVs at very low doses in the mouse heart following intravenous injection of sEV-L1-Zr. Additionally, *ex vivo* experiments using a Langendorff rat heart perfusion model confirmed targeted accumulation of the vesicles in cardiomyocytes as compared to other cells in the heart compartment. This approach provides a promising platform for sensitive and stable in *vivo* tracking of sEVs, advancing their application in both diagnostic imaging and regenerative therapies.

## INTRODUCTION

sEVs are a heterogeneous group of nanosized vesicles secreted by nearly all cell types. They carry a diverse array of biomolecules, including genetic material, proteins, and lipids, which are incorporated into their lumen or attached to their surface during biogenesis [1–6]. These vesicles play a crucial role in intercellular communication, acting as key regulators in both physiological and pathological processes [7, 8]. By transferring their molecular cargo, sEVs can sequentially modulate the behaviour of recipient cells, making them promising tools for diagnosis and therapy [9–13]. This potential is particularly evident in the context of cardiovascular diseases such as myocardial ischemia without infarction, various forms of angina, acute coronary syndrome, and others [13, 14]. Currently, numerous observational and interventional clinical trials are exploring the applications of sEVs in the cardiovascular field, highlighting their growing significance in clinical research [13].

For diagnostics and therapies, it is critical to develop imaging platforms that allow monitoring the biodistribution of sEVs [15]. The main available imaging techniques are based on fluorescence, luminescence, positron emission tomography (PET)/magnetic resonance (MR), and single photon emission computed tomography (SPECT) [15]. For *in vivo* imaging, methods that use PET/MRI or SPECT/computed tomography, both requiring radionuclide labeling, offer greater sensitivity and absolute quantification, allowing the acquisition of images with anatomical details [15]. The most common radioisotopes used for PET imaging are ^18^F, ^68^Ga, ^86^Y, ^64^Cu, ^82^Rb, and ^89^Zr — with half-lives of 109.7 min, 67.7 min, 14.7 h, 12.7 h, 1.3 min, and 78.4 h, respectively [16]. In a previous study, we developed a strategy to radiolabel sEVs with ^64^CuCl_2_ for PET/MRI imaging. Radiolabeled sEVs were detected in organs with low accumulation, such as the brain (0.4–0.5% ID g^−1^), with further anatomical brain location determined by MRI [17]. Unfortunately, the preparation of ^64^CuCl_2_ is expensive, complex and has a short half-life, which makes it difficult for longer monitoring. Other studies have used ^86^Y, which has a similar half-life to ^64^Cu. Besides the same ^64^Cu limitation to be used in longer sEV biodistribution studies, it also adds the limitations of sub-optimal decay characteristics and difficult radionuclide production and purification protocols [18]. Isotopes like ^111^In and ^67^Ga also exhibit favourable physical half-lives and behaviour for sEV-based applications; however, the inherent limitations of SPECT over PET imaging platforms make them less attractive. PET imaging is better due to its emission of positrons, rather than single photons, enabling higher-resolution and quantitative imaging [18, 19]. ^99m^Tc also suffers from the limitations of SPECT imaging and has a half-life as too short for imaging. In contrast, ^124^I manifested a near-ideal half-life for imaging; however, such as the high cost of the isotope, its relatively low resolution due to the high energy of its positrons, and the significant dehalogenation of ^124^I-labeled sEVs *in vivo* collectively limit its ultimate clinical potential [20, 21].

The physical properties of ^89^Zr make it highly advantageous for sEV-based imaging applications. With a half-life of approximately 78.4 h, low positron energy of 395.5 keV, and favourable characteristics such as safety, ease of production, and *in vivo* stability, ^89^Zr offers several benefits for imaging studies [22–25]. A critical aspect of designing effective PET tracers involves forming stable radiometal chelations. DFO is the most common chelator used for ^89^Zr PET imaging, forming complexes with Zr^4+^ at room temperature [26]. However, DFO’s stability remains a concern; in several preclinical studies, leaching of ^89^Zr^4+^ from the DFO complex has been observed as early as one-hour post-administration [23, 26–28]. This instability poses significant challenges when detecting sEVs at low concentrations, especially below 1% of the initial signal. Recent research indicates that 1,4,7,10-tetraazacyclododecane-1,4,7,10-tetraacetic acid (DOTA) forms more stable complexes with Zr^4+^[26], but this typically requires high temperatures (∼90°C) for optimal chelation. Such elevated temperatures are incompatible with the structural integrity of sEVs. Therefore, there is a pressing need to develop chelators capable of rapidly forming stable complexes with Zr^4+^ at room temperature, ensuring effective and reliable imaging of sEVs.

Here, we present the development of a novel PET labeling kit for sEVs utilizing ^89^Zr. This kit is designed to enable *in vivo* tracking of sEVs for applications in diagnostics and therapeutics, with a particular focus on cardiac health. sEVs were labeled with ^89^Zr^4+^ using an innovative chelator, designated L1 (**Scheme 1**). The L1 chelator is a hybrid construct combining features of DOTA and DFO: it replaces the three acetate arms of DOTA with hydroxamate groups similar to those in DFO, while retaining one acetate arm for conjugation to sEVs. The hydroxamate functionalities in L1 enhance the kinetics of ^89^Zr^4+^ binding at room temperature, and the macrocyclic structure confers superior stability compared to DFO alone. Following labeling, the ^89^Zr -labeled sEVs were administered to mice, and their biodistribution and accumulation were monitored over 24 h using combined PET/MRI imaging. Our results demonstrated that this imaging approach can detect sEVs with high sensitivity, identifying as little as 1% of injected sEVs within the heart tissue after 24 h. *Ex vivo* studies utilizing a Langendorff-perfused heart model further confirmed that ^89^Zr-labeled sEVs preferentially accumulated in cardiomyocytes. Overall, the proposed sEV labeling strategy offers several advantages over existing methods, including enhanced sensitivity, improved stability, superior spatial resolution, and cost-effectiveness.

## RESULTS

### Synthesis of a new ^89^Zr^4+^ chelator

To synthesize a novel chelate capable of complexing Zr⁴⁺ at room temperature, we initially prepared the Arm *N*-(benzyloxy)-2-bromo-*N*-methylacetamide (**Fig. 1a**). This arm introduces the hydroxamic acid functional group into the final chelate, which is essential for promoting effective complexation with Zr⁴⁺ under ambient conditions. We initiate the synthesis of the Arm using N-methylhydroxylamine. In brief, the amine group of the N-methylhydroxylamine was protected with Di-tert-butyl decarbonate (BOC) before the substitution reaction with benzoyl chloride, which resulted in **A2 (Fig. S1)**. Following the deprotection of BOC (**A3; Fig. S1**), the **Arm** was obtained by incorporating bromoacetyl bromide (**Fig. S1**). After having the hydroxamic acid pendant arm, **L1** was synthesized from 1,4,7,10-tetraazacyclododecane-1-benzyl acetate (**C3**) in a two-step chemical reaction (**Fig. 1b**). Briefly, **C4** was prepared by first the deprotection of the -BOC of the **C3,** synthesized from Cyclen following standard protocols [29], using TFA and followed by the addition of **Arm** to the deprotected **C3 (Fig. S2)**. After workup, the residue was purified via column chromatography to afford a pure solid. Deprotection of the benzyl group of **C4** was carried out selectively. To achieve a full deprotection, the reaction was performed at 70°C temperature for 48-72 h. The crude oil-like product was dissolved in methanol and precipitated with diethyl ether to quantitatively yield L1 as a white fluffy solid (**Fig. S3**).

**Figure 1.**
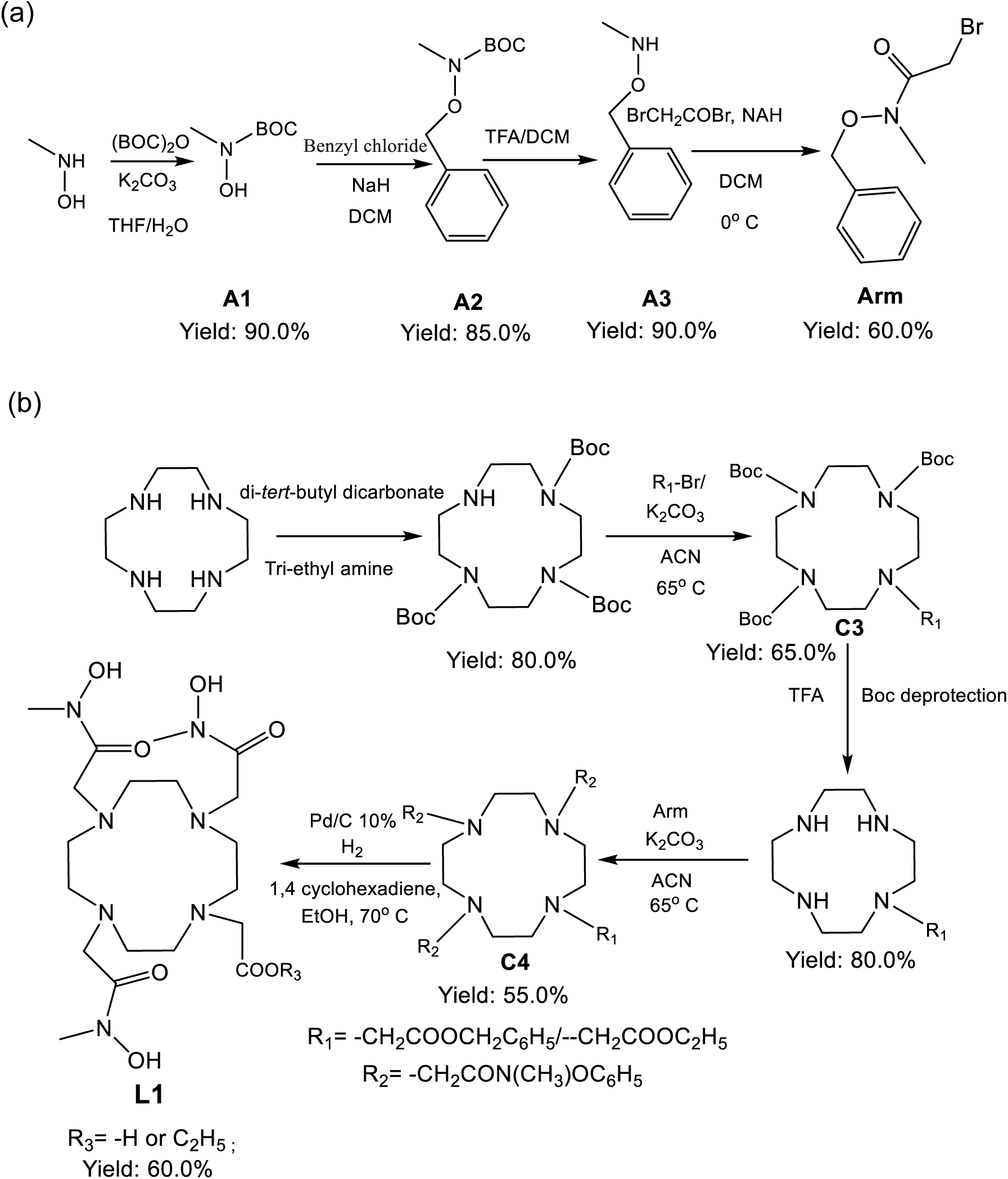
Synthesis of the chelator L1. (a) Synthesis of the Arm (*N*-(benzyloxy)-2-bromo-*N*-methylacetamide) (b) and L1.

To demonstrate the ability of the new chelator to complex Zr⁴⁺, we incubated L1 (1 mM) with an equimolar concentration of Zr⁴⁺ at room temperature and monitored the complex formation using HPLC (**Figs. 2a and 2b**). Initially, we assessed the complexation of Zr⁴⁺ by known chelators such as DFO and DOTA by HPLC (**Figs. S4a-S4b**). Upon incubation with Zr⁴⁺ at room temperature, the new chelator–Zr⁴⁺ complex eluted alongside the salts from the buffer, while the peak corresponding to the uncomplexed chelator diminished in intensity. Based on the decrease of the peak of the chelator in the HPLC run, our results indicated that approximately 65% of the initial DFO was complexed with Zr⁴⁺, whereas DOTA showed negligible complexation under these conditions (**Figs. S4a-S4b**). We tracked the formation of L1-Zr complexes by monitoring the reduction in the area of the free L1 peak across different buffer systems (**Fig. 2c**). In PBS buffer, the overall complexation yield was approximately 25%, whereas in Tris-HCl buffer, it reached about 57%, which is slightly lower than the complexation efficiency observed with DFO alone. Furthermore, the ^1^H NMR spectrum of L1-Zr complex exhibits a clear downfield shift in the –CH₃ peaks (3.21 ppm to 3.47 ppm) of the hydroxamate arm and the –CH₂ peaks (2.9 ppm- 3.5 ppm) of the cyclam rings compared to free L1, indicating the formation of the L1-Zr complex (**Fig. S4c**). Similarly, -C=O group in the hydroxamate arm has a clear shift from 1560-1700 to 1590-1750 cm^-1^ in FTIR due to the formation of the L1-Zr (**Fig. S4d**). Further O–H stretching band of the hydroxamate moiety appears at 3850 cm⁻¹ (L1-Zr), attributable to the formation of a strong Zr–OH bond that weakens the hydrogen bonding of the free –OH group.

**Figure 2.**
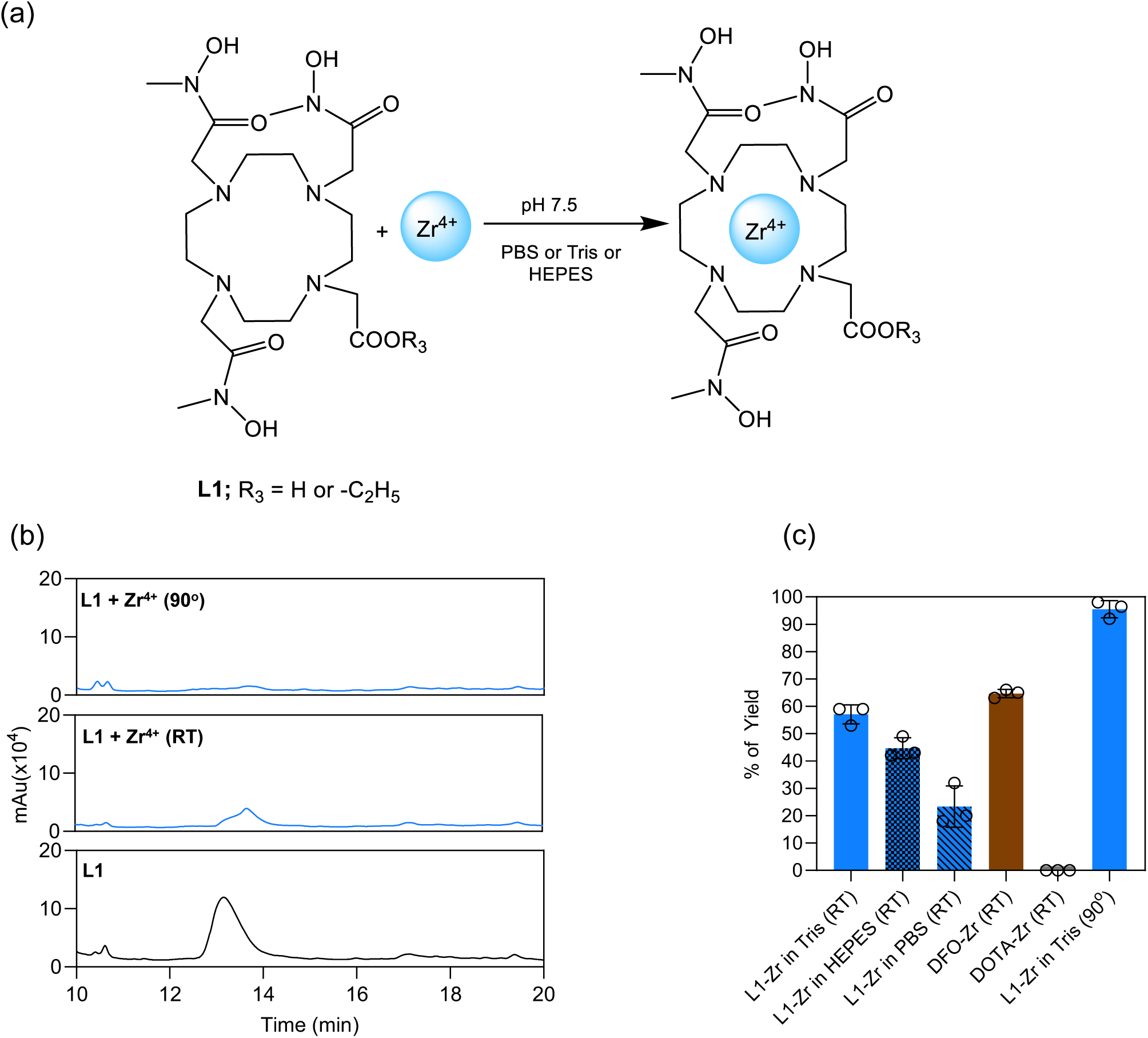
**Complexation of Zr with L1 chelator**. (a) Scheme of the complexation reaction. (b) Representative HPLC analyses of free L1 at room temperature, L1 mixed with Zr^4+^ at room temperature (RT) after 1 h and L1 mixed with Zr^4+^ and refluxed at 90°C after 1 h. (c) Yield data for the complexation of Zr^4+^ with the L1 chelator under various experimental conditions and compared with other chelators (DFO and DOTA; complexation reaction for 1 h. Results are Mean ± SD (n=3 independent experiments).

Overall, we have successfully synthesized a novel chelator for Zr⁴⁺ through a multi-step experimental process, resulting in a milligram-scale yield. The new chelator demonstrates comparable complexation efficiency to DFO under ambient conditions, following optimization of the reaction buffer.

### Surface labeling of sEVs with ^89^Zr^4+^

To demonstrate that our novel chelator can effectively label sEVs with Zr⁴⁺, we initially selected two sEV sources, human umbilical cord blood mononuclear cells (hUCB-MNCs) and human urine, to evaluate the robustness of our labeling strategy. Both sEV types were thoroughly characterized according to the latest ISEV guidelines [30], assessing parameters such as size (**Fig. S5a**), purity (**Fig. S5b**), surface charge (zeta potential; **Fig. S5c**), morphology, and surface protein expression. Despite their distinct origins, the sEVs displayed similar size distributions, with some variations in zeta potential and particle count per microgram of protein (**Fig. S5a–c**). Notably, a more negative zeta potential was associated with a lower number of particles per microgram of protein. Transmission electron microscopy (TEM) confirmed the characteristic cup-shaped morphology of both sEV sources, with no significant aggregation observed (**Fig. S5d**). Western blot analyses further validated the presence of key sEV markers across samples from two different donors for each source, demonstrating consistency and reliability in our characterization approach (**Fig. S5e**). To verify sample purity, we tested for proteins absent in sEVs- Apolipoprotein A1 (ApoA1) in hUCB-MNC-derived sEVs and Tamm-Horsfall protein (THP) in urine-derived sEVs- and found negligible contamination in both preparations. Additionally, hallmark sEV membrane proteins, including tetraspanins CD9 and CD63, as well as Alix (only for urine EVs), involved in vesicle biogenesis and release, were detected in the particles.

Next, we assessed the conjugation of L1 with sEVs derived from urine, leveraging the free amine groups present on the sEV surface (**Fig. 3a and S6a**). The efficiency of the reaction was evaluated by measuring the remaining free amine groups using a fluorescamine assay (**Fig. S6b**). A decrease in free amines post-labeling indicated successful conjugation of L1 to the sEVs. Approximately 44% of the available amine groups reacted with L1 at pH 7.5 (**Fig. S6c**). This means that each sEV has approximately 16,000 conjugated L1 molecules. Subsequently, EV-L1 were complexed with Zr^4+^ and purified with SEC columns (**Fig. 3a**). Size analysis showed no significant changes in sEV dimensions (**Fig. 3b**). As a control, a separate batch of sEVs was first conjugated with DFO via surface amine groups, then complexed with Zr^4+^. Similar to the L1-conjugated sEVs, these exhibited no significant size alterations (**Fig. 3b**). The slight increase in zeta potential following Zr^4+^ labelling of the sEVs suggests successful complexation and formation of EV-L1-Zr complexes, as both L1-Zr and DFO-Zr complexes carry a +1 charge (**Fig. 3c**). Interestingly, no significant differences in zeta potential were observed between EV-DFO-Zr and EV-L1-Zr, indicating comparable surface charge characteristics.

**Figure 3.**
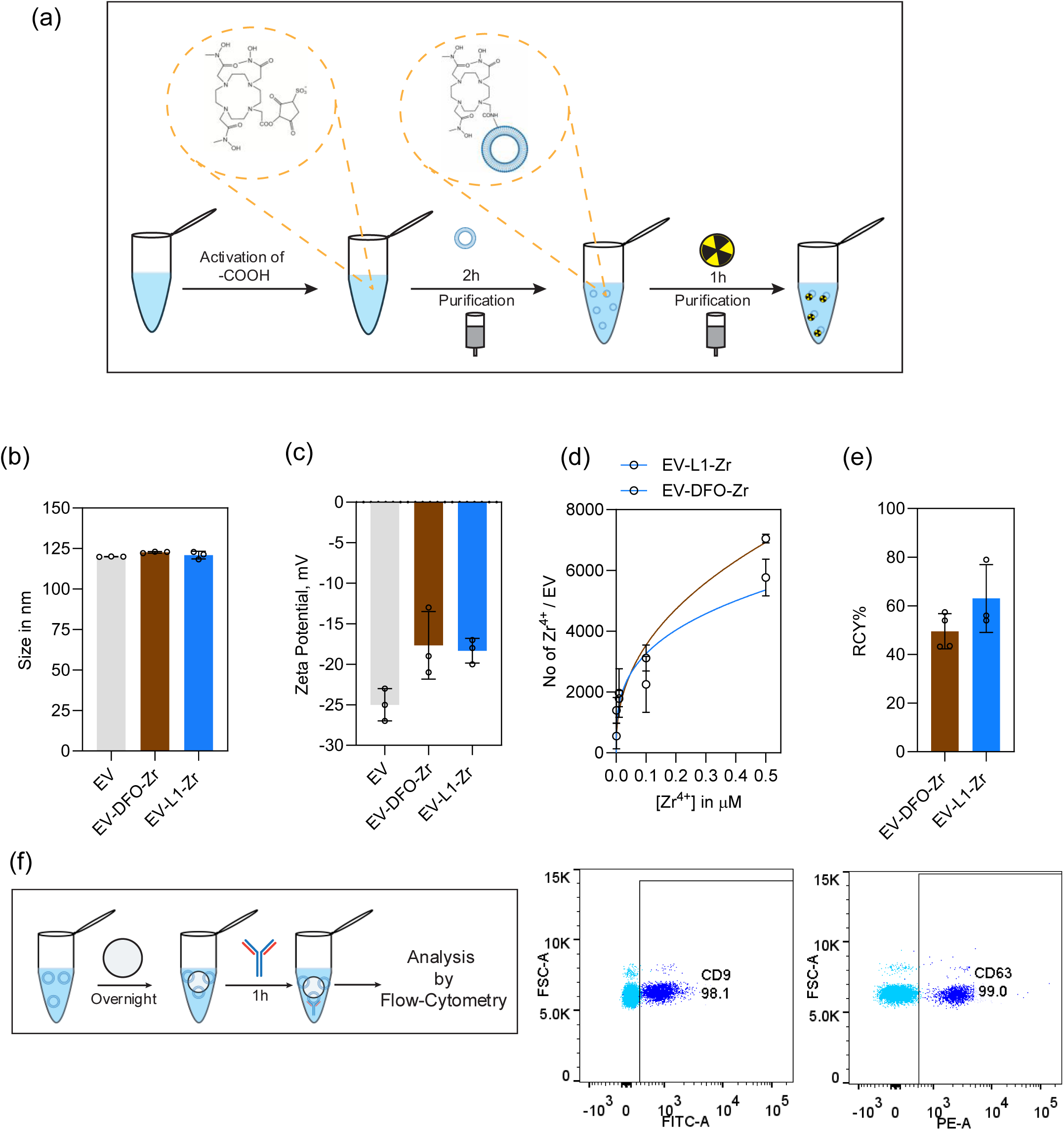
Radiolabeling and characterization of the Zr-labelled sEVs. (a) Schematic presentation of the radiolabeling process. (b) Quantification of sEV size before and after Zr^4+^ labeling. (c) Quantification of zeta potential of sEVs before and after Zr^4+^ labeling. (d) Titration of EV-L1 or EV-DFO with various concentrations of Zr^4+^. (e) Radiochemical yield (RCY) of the labelling process. This was calculated by the ratio between the amount of activity present after the labelling versus the amount of activity used for the labelling. In b-e, results are Mean ± SD (n=3 independent runs). (f) Flow cytometry quantification of sEV markers, including CD63 and CD9, on Zr-labelled sEVs. Effect of the labelling process on the surface markers of sEVs.

To determine the maximum number of Zr⁴⁺ ions that can be incorporated into EV-L1, titration experiments were conducted using varying concentrations of ZrCl₄ (**Fig. 3d**). The results show that EV-L1-Zr can incorporate approximately 5,770 Zr^4+^ ions, and thus a complexation efficiency of 36%, when complexation is performed in Tris-HCl buffer conditions. This value is slightly lower than the approximately 7,050 Zr^4+^ ions per EV achieved with EV-DFO-Zr, with a complexation efficiency of 44%. Most importantly, the native sEVs can absorbed maximum 149 ± 50 Zr^4+^ per sEV. This results clearly indicate the labelling of sEVs with Zr^4+^ was done due to the complexation of the L1- Zr^4+^ and DFO- Zr^4+^. Using this optimized approach, we proceeded to radiolabel the sEVs with ^89^Zr^4+^. After each step, purification was carried out using SEC columns, which effectively removed unreacted chelator and free ^89^Zr^4+^ from the mixture. The final radiochemical yield was approximately 60%, comparable to that of EV-DFO-Zr (**Fig. 3e**). Importantly, this labeling strategy did not compromise the integrity of EV surface markers (**Fig. 3f**).

Altogether, we successfully established a two-step labelling method for labelling sEVs with Zr⁴⁺ using the novel chelator L1. The radiochemical yield achieved with L1 was comparable to that of DFO.

### Stability and biological activity of sEVs labeled with ^89^Zr^4+^

The radiochemical purity of EV-L1-^89^Zr and EV-DFO-^89^Zr after purification by SEC column was characterized by instant thin-layer chromatography (ITLC). The free ^89^Zr moved along the TLC strip, whereas EV-L1-^89^Zr stayed at the bottom of the strip (**Fig. 4a**). Our results showed that the purification was successful and a purity of approximately 100% was obtained for both complexes.

**Figure 4.**
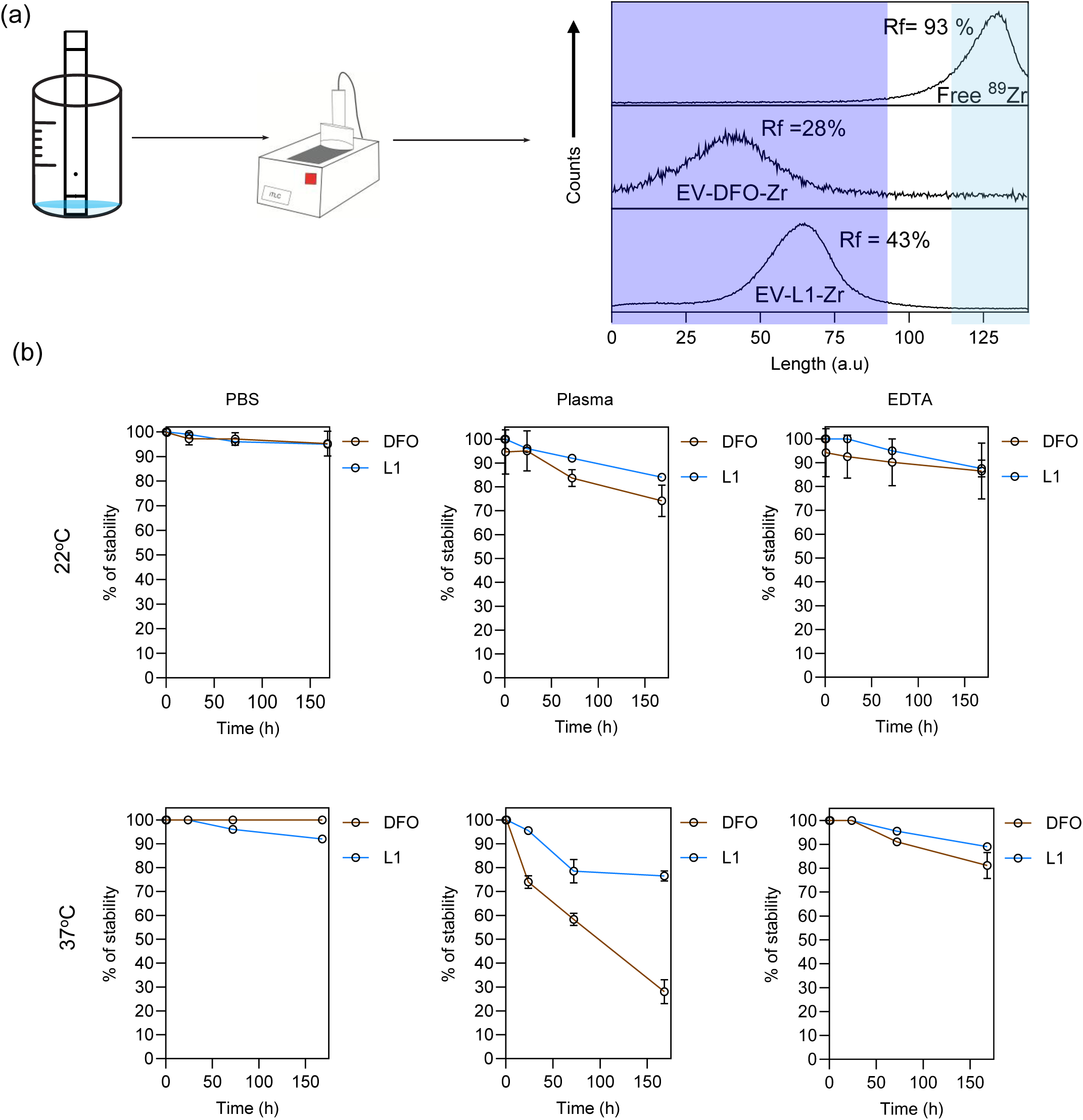
Purity and stability measurement on Zr-labelled sEVs. (a) Purity of EV-DFO-Zr and EV-L1-Zr was assessed using ITLC. Free ZrCl_4_ migrates with the solvent front, while EV- DFO-Zr and EV-L1-Zr remain at the origin of the TLC strip. These results demonstrate that the purification process yields products with 100% purity. (b) Stability measurements of EV-DFO-Zr and EV-L1-Zr in different buffers like PBS, plasma (50% diluted using dilution buffer containing EDTA and PBS), and 1M EDTA at 22 °C and 37 °C for 7 days. Results presented are Mean ± SD (n=3 independent experiments).

For *in vivo* applications, it is very important that the ^89^Zr labelling is stable and the ligand does not leach overtime from the chelator. To evaluate the stability, EV-L1-^89^Zr and EV-DFO-^89^Zr (∼ 1× 10^10^/mL) were incubated over a period of 7 days in various solutions, including PBS, human plasma (50% diluted with a buffer containing ethylenediamine-*N*,*N*,*N’*,*N’*-tetraacetic acid (EDTA) and PBS), and 1 M EDTA, at both 22°C and 37°C (**Fig. 4b**). At 22°C, the results demonstrated that EV-L1-^89^Zr exhibited greater stability compared to EV-DFO-^89^Zr across different time points (**Fig. 4b**). Both complexes showed reduced stability in human plasma relative to PBS and EDTA solutions. At 37°C, stability declined in plasma for both complexes, with EV-DFO-^89^Zr retaining only about 75% stability after one day, whereas EV-L1-^89^Zr maintained over 95% stability (**Fig. 4b**).

Next, we assessed whether the labeling procedure impacted the biological properties of sEVs. First, we examined the cytotoxicity of EV-L1-Zr and native sEVs (both at 1×10^9^ particles/mL) on human endothelial cells following 48 h of exposure (**Fig. 5a** and **Fig. S7a** for EV-DFO-Zr). After incubation, cells were washed, and cell numbers were quantified using Hoechst staining. As a control, we also evaluated the cytotoxic effects of 10 nM Zr^4+^, the concentration of Zr that we estimate to be present on EV-L1-Zr. Our results showed no significant differences in the number of live cells among the control, cells treated with sEVs, EV-L1-Zr or Zr^4+^ (**Fig. 5b** and **Fig. S7b** for EV-DFO-Zr). Second, we investigated whether Zr-labeling affected EV internalization in human endothelial cells (**Fig. 5c** and **Fig. S7c** for EV-DFO-Zr). For this, EVs and EV-L1-Zr were labeled with an intraluminal fluorescent dye (CFSE), then incubated with cells for 4 h. Following incubation, cells were washed to remove non-internalized sEVs and analyzed via confocal microscopy. We observed no significant difference in the percentage of cells labeled with native sEVs versus EV-L1-Zr. Third, we evaluated the bioactivity of sEVs (isolated from hUCB-MNCs), EV-L1-Zr, and EV-DFO-Zr on human endothelial cells cultured under hypoxic conditions. Previous studies from us have demonstrated that MNC-derived EVs promote survival in these cells [17, 31, 32]. Cells were subjected to ischemic conditions (glucose-free media in the absence of oxygen) for 2 h, then treated with EVs, EV-L1-Zr (4.5 × 10^9^ particles/mL), or EV-DFO-Zr (4.5 × 10^9^ particles/mL) for 48 h under normoxic conditions with complete media (**Fig. 5d** and **Fig. S7d** for EV-DFO-Zr). The results indicated no significant difference in bioactivity between EVs and EV-L1-Zr, suggesting that our labeling process does not compromise EV function. Similar findings were observed with EV-DFO-Zr (**Fig. S7a-d**).

**Figure 5.**
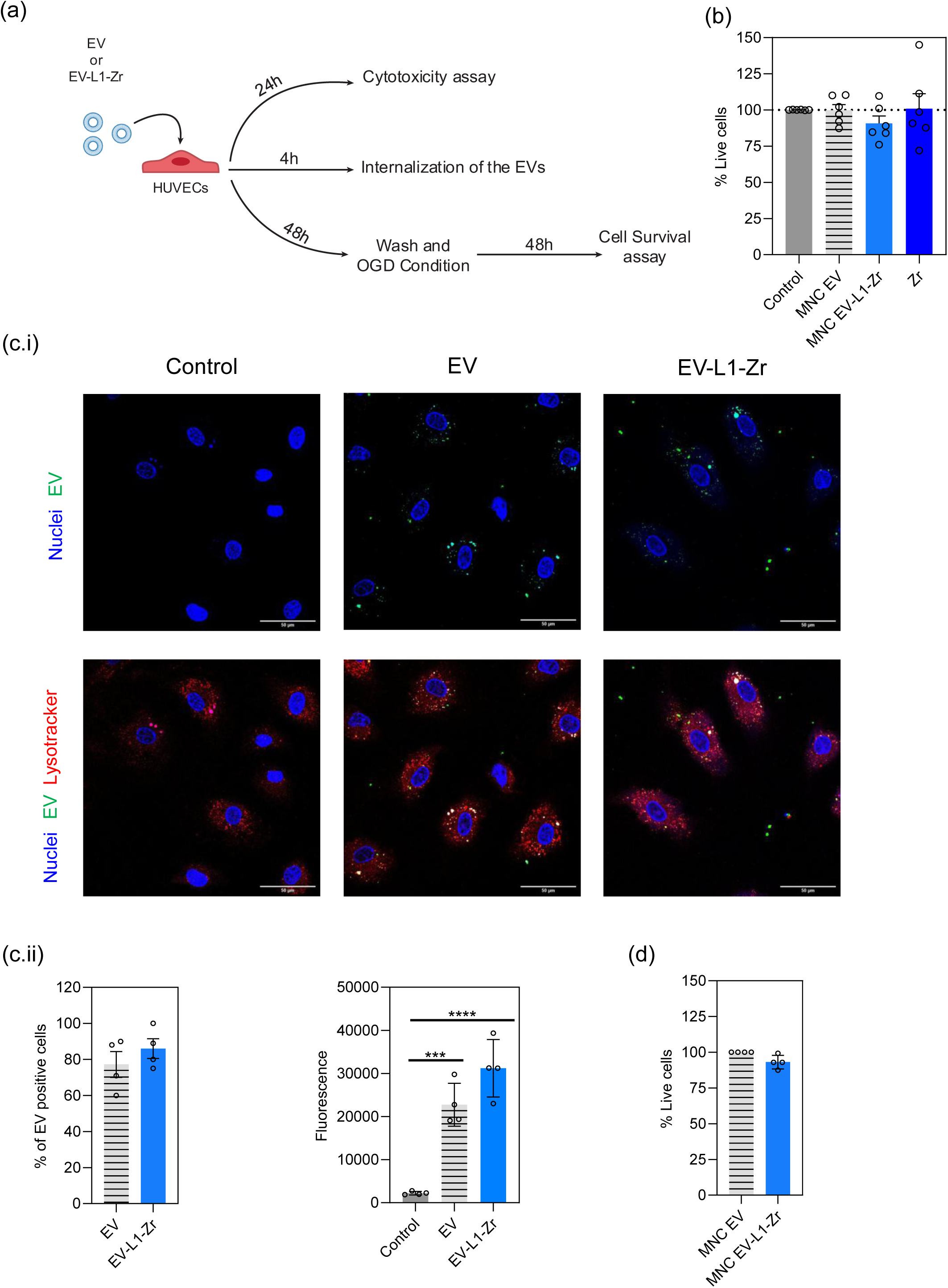
Cytotoxicity, cellular internalization, and bioactivity of Zr-labelled sEVs. (a) Experimental setup. (b) Cytotoxicity measurements of EV, EV-L1-Zr, and Zr (10 nM) on HUVECs. (c) Internalization of sEV and EV-L1-Zr on HUVECs. (d) Bioactivity of EV-L1-Zr (see Materials and Methods). The percentage of cell survival values resultant from the experiment, which indicates the pro-survival effect. In b-d, results are Mean ± SD (n=4-6 independent experiments, each experiment has 3 technical replicates).

Overall, our data demonstrate that EV-L1-Zr are more stable in plasma at 22°C and 37°C compared to EV-DFO-Zr. Importantly, the labeling strategy did not alter the cytotoxicity profile, cellular internalization, or bioactivity of EV-L1-Zr.

### *In vivo* biodistribution and *ex vivo* heart cellular accumulation of sEVs labelled with ^89^Zr^4+^

The biodistribution of EVs was monitored over a 24 h period using EV-L1-Zr (∼160–220 μCi per 2.5×10^10^–3.5×10^10^ particles of sEVs which is much lower than the 600 μCi injected for ^64^Cu^2+^ in a previous study performed by us [17]) via PET/MRI imaging following intravenous injection into the tail vein of C57BL/6J mice (**Fig. 6a**). For these experiments, EV-L1-Zr (n=3 mice) was imaged by PET-MRI at 1, 5, and 24 h post-administration (**Fig. 6b**). All PET scans were performed using a prototype of a high-acceptance small-animal PET based on resistive plate chambers (RPC-PET)[33]. The whole body of a mouse was scanned in three sections (liver-heart, brain, kidney-bladder). Images were reconstructed using OSEM algorithm and a cubic voxel of 0.5 mm width. Height fiducial markers that can be viewed both in the PET and MRI imaging were placed in the mouse bed, and MRI imaging was done after PET without moving the mice from the bed, under anesthesia. The highest accumulation of sEVs was observed in the liver and spleen. At 24 h, animals were sacrificed, and organ activity was quantified using a scintillation counter (**Fig. 6c**). Approximately 50% of the sEVs localized in the liver, with around 20% in the spleen. Notably, there was very low activity in the blood at 24 h (0.2 %ID/g), indicating almost complete delivery of the sEVs to the organs. Due to the long half-life and low positron emission energy of ^89^Zr, sEVs were readily detectable in the heart (average 0.32 **Figs. 6b and 6c**). Importantly, the L1 chelator conferred stability and minimized ^89^Zr leaching from sEVs. EV-L1-Zr showed approximately 1.5% accumulation in bone tissue (**Fig. 6d**), compared to about 6% for EV-DFO-Zr (**Fig. 6d**) and approximately 10% for EV labelled with ^89^Zr-(Oxinate) [34]. These findings demonstrate the superiority of our novel chelator and labelling approach for long-term tracking of sEVs, particularly in organs with low accumulation. Our results further show that the heart with accumulation of sEVs over time (**Fig. 6e**).

**Figure 6.**
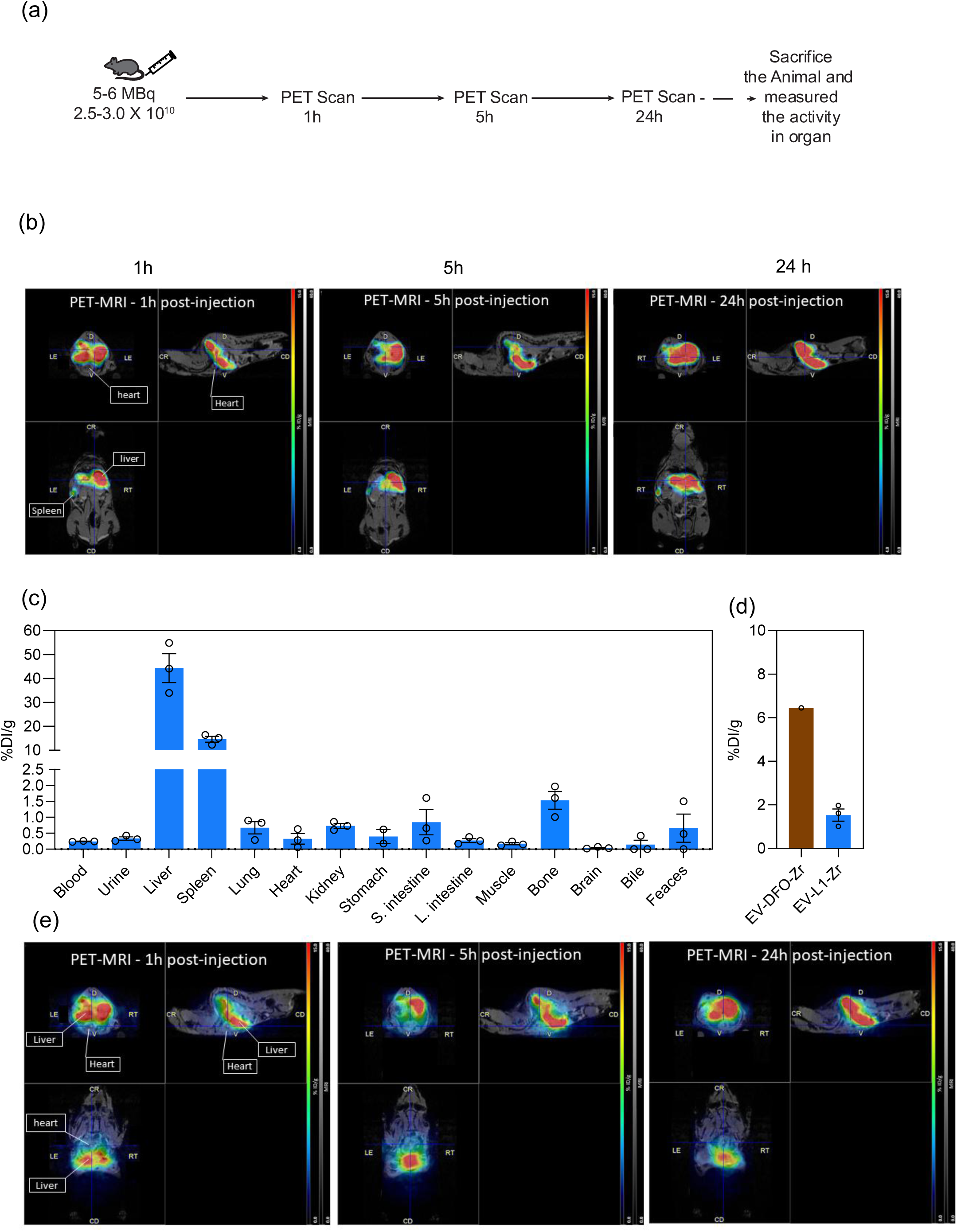
*In vivo* biodistribution of Zr-labelled sEVs. (a) Schematic presentation of the experimental design. (b) PET/MRI images of the EV-L1-Zr in different time points after 1 h, 5 h, and 24 h of the injection. (c) Biodistribution of EVs at 24 h post-administration using a well counter. (d) Accumulation of free Zr^4+^ in the bone. (e) Accumulation of EVs in the heart after 24 h using PET/MRI. The bars represent mean ± SD (n=3 animals).

To evaluate the cellular distribution of sEVs within the heart, we employed an *ex vivo* Langendorff heart perfusion (LHP) model using hearts harvested from 10-week-old Wistar Han rats. Since our LHP setup could not accommodate radioactive labelling, we labelled the sEVs with non-radioactive Zr^4+^. In this protocol, hearts were perfused with EV-L1-Zr (7.5 × 10^9^ particles suspended in 30 mL of Krebs buffer, thus 2.5 × 10^8^ EVs/mL) for one hour (**Fig. 7a**). After perfusion, the system was washed, and cardiomyocytes along with non-cardiomyocyte cells were isolated from the hearts using a protocol published elsewhere[35]. The cardiomyocyte cells were characterized by western blot using Troponin (as cardiomyocyte marker) and Vimentin (as non-cardiomyocyte marker) (**Fig. S8**). These cells were counted, freeze-dried, and the quantity of Zr^4+^ within each cell population was measured using Inductively Coupled Plasma Mass Spectrometry (ICP-MS). Our data revealed a higher accumulation of sEVs in cardiomyocytes ∼60% compared to non-cardiomyocytes ∼40% (**Fig. 7b**). To corroborate these findings, we performed another experiment using fluorescence-labeled EVs (EV-L1-Zr-Dil) and analyzed their localization within the heart tissue via confocal microscopy. Post-Langendorff perfusion, hearts were snap-frozen in optimal cutting temperature (OCT) compound, sectioned transversely, and subjected to immunohistochemistry. Confocal imaging confirmed that sEVs predominantly accumulated in cardiomyocytes (**Figs. 7c-7d**). This complementary approach further supports our conclusion that sEVs preferentially localize within cardiomyocytes in the heart.

**Figure 7.**
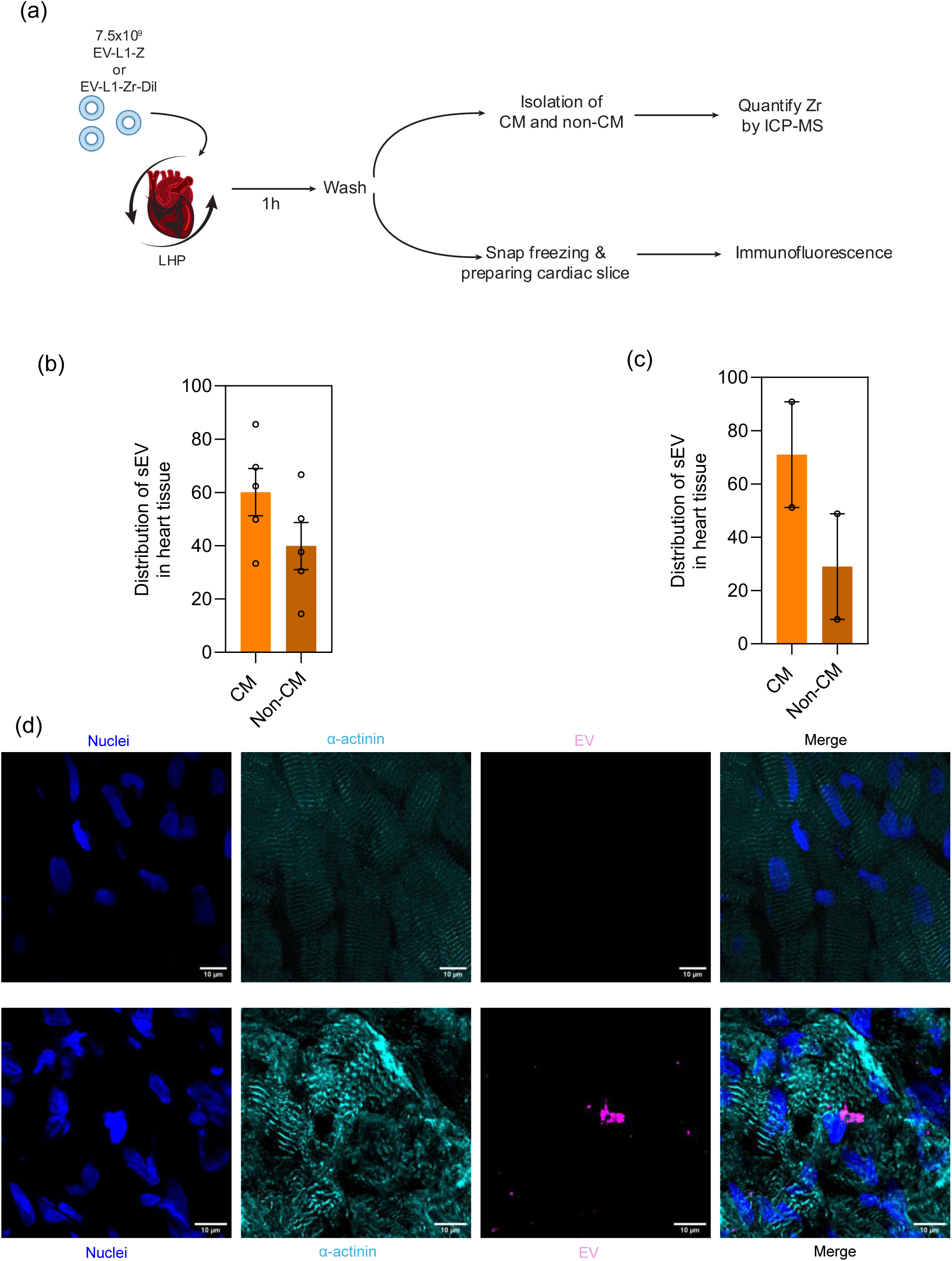
*Ex vivo* biodistribution of Zr-labelled EVs on a Langendorff perfusion model. (a) Schematic presentation of the experimental design. (b) Distribution of sEVs in heart cells, as assessed by ICP-MS analysis following their extraction from cardiac tissue. (c) Distribution of sEVs in heart cells as evaluated by confocal microscopy. (d) Representative confocal images showing the accumulation of sEVs in CMs. Results in b and c are Mean ± SD (n=2-5 rat hearts).

Overall, EV-L1-Zr exhibited high *in vivo* stability with PET/MRI showing predominant sEV accumulation in the liver and spleen at 24 h. *Ex vivo* Langendorff perfusion revealed higher uptake in cardiomyocytes than non-cardiomyocytes, confirmed by both ICP-MS and confocal microscopy analyses. These results demonstrate the robustness of the L1 chelation strategy for long-term *in vivo* imaging.

## DISCUSSION

We have developed a novel radiolabeling strategy that enhances PET/MRI imaging capabilities. This approach leverages a newly designed ligand that can be immobilized on the surface of sEVs, enabling efficient complexation with the radionuclide ^89^Zr^4+^ at room temperature. Compared to existing radiolabeling methods, our strategy offers several key advantages: it allows for extended tracking of sEVs over longer periods and utilizes a relatively cost-effective radionuclide. In our experiments, we used activities ranging from 160 to 220 μCi per 2.5×10^10^ to 3.5×10^10^ sEV particles. This activity can be scaled up to milliCurie levels per 1×10^9^ sEV particles by employing higher concentrations of ^89^Zr^4+^. Our findings demonstrate that EV-L1-Zr exhibits greater stability in plasma compared to EV-DFO-Zr. Importantly, these labeled sEVs do not induce cytotoxicity and do not exhibit altered cellular internalization or pro-survival activity. *In vivo*, EV-L1-Zr can be monitored via PET/MRI for up to 24 h, enabling tracking of sEV accumulation, including in the heart.

One of the primary challenges associated with using ^89^Zr^4+^ is ensuring the stability of its labeling process under *in vivo* conditions. Biodistribution studies in mice have shown that approximately 5–10% of injected ^89^Zr^4+^-DFO-labeled antibodies accumulate in the bones, indicating *in vivo* dissociation of the radiometal from the chelator DFO [36, 37]. The commonly used chelator DFO forms complexes with ^89^Zr^4+^, but these are often unstable because DFO can only satisfy six of the eight possible coordination sites, leaving two sites uncoordinated and vulnerable to dissociation. Efforts to enhance DFO’s chelating ability by synthesizing derivatives such as DFO* or DFOCyclo* aimed to provide additional coordination sites but with limited success [37]. In contrast, macrocyclic ligands like DOTA form more stable complexes with ^89^Zr^4+^. Structural studies of DOTA-Zr complexes have revealed an octa-coordinated configuration involving macrocyclic nitrogen atoms and acetate pendant arms, which explains the higher stability of these complexes [26]. However, the complexation kinetics with DOTA are slow and typically require elevated temperatures. In our study, we have developed a novel ligand designed to satisfy all eight coordination sites of ^89^Zr^4+^ under room temperature conditions, resulting in a highly stable complex. This chelator combines elements of DOTA and DFO: four arms from DOTA, with three arms replaced by hydroxamate groups similar to DFO, and the remaining coordination site provided by a nitrogen donor from the macrocyclic ring. The hydroxamate arms facilitate rapid complexation kinetics at room temperature, while the macrocyclic component enhances the overall stability of the ^89^Zr^4+^ complex.. This design ensures that the chelator can effectively coordinate all eight sites of ^89^Zr^4+^, resulting in a more stable and kinetically favorable complex suitable for *in vivo* applications.

The present study advances both the design of a novel chelator for ^89^Zr^4+^and the development of an effective complexation methodology. Our findings highlight the critical influence of buffer pH on the kinetics of complex formation with the ligand L1. Specifically, complexation in PBS buffer was notably slow, whereas in Tris-HCl buffer, the kinetics closely resembled those observed with deferoxamine (DFO) (**Fig. S2c**). This difference is primarily due to the strong affinity of phosphate groups toward Zr^4+^, forming stable complexes [38, 39], which impedes the reaction rate at room temperature and results in lower yields of L1-Zr conjugates in PBS. To optimize radiolabeling efficiency, we performed the process in Tris-HCl buffer at pH 7.5, subsequently switching to PBS (pH 7.5) during the purification step. Regarding radiolabeling agents, although the most commonly used form of ^89^Zr for such applications is ^89^Zr(Ox)_2_, which is highly stable even at low pH but toxic [26, 40], we utilized ^89^ZrCl_4_ in our study. Neutral ^89^ZrCl_4_ tends to form hydroxo- or oxo-bridged species in aqueous buffers. Importantly, in the absence of oxalate ligands, ^89^ZrCl_4_ favors complexation with tetraazamacrocyclic chelators over the formation of ^89^ZrOH species, thereby enhancing labeling efficiency [41]. Our observations also indicate that the radiochemical yield of sEV labeling is influenced by the dilution of the reaction mixture. For ^89^ZrCl_4_, excess solvent can be readily removed via evaporation, whereas similar treatment with ^89^Zr(OX)_2_ results in the precipitation of insoluble radionuclides, leading to activity loss and reduced overall efficiency. Notably, the maximum number of zirconium atoms incorporated per EV was approximately 5,770, which is twenty times higher than our previous study using ^64^Cu for EV radiolabeling [17]. This improvement significantly enhances the sensitivity and enables longer-term monitoring. Furthermore, the disparity in labeling efficiency appears to stem from the higher availability of free amino (-NH_2_) groups compared to free thiol (-SH) groups per EV, serving as a primary factor influencing conjugation success.

Previous studies have shown the labeling of sEVs with ^89^Zr^4+^ using DFO as a chelator [42–44] or the direct intraluminal radiolabeling using [^89^Zr]Zr(oxinate)_4_ [34]_;_ however, all those strategies have limitations (**Supplemental Table 1**). First, they showed lower stability than our current strategy. Using our labelling strategy, we were able to show that the *in vivo* leaching of Zr and subsequent accumulation in bone is negligible (∼1.5 %). This was ∼7-fold and ∼4-fold lower, respectively, than previously reported ^89^Zr-Oxinate (∼10%)[34] and EV-DFO-Zr (∼6%; current results). Second, the yield of labeling in previous radiolabeling strategies was relatively inferior to our strategy (**Supplemental Table 1**).

In recent years, EVs from different sources have been used in the treatment of cardiovascular diseases[13]. Unfortunately, in most studies, it is relatively unknown which cardiac cells internalize sEVs when a therapeutic strategy is reported. To identify the cellular compartment in the heart that accumulates higher levels of sEVs, we performed the Langendorff perfusion heart model (**Fig. 7a and 7b**). The results from the ICP-MS confirm that the majority of the sEVs were taken by the cardiomyocytes (**Fig. 7b**). To our knowledge, this is the first direct demonstration of sEV delivery to cardiomyocytes using an anatomical setup. This result was further supported by confocal microscopy images using sEV-L1-Zr-Dil where the majority of the sEVs were colocalized with the CM (**Fig. 7c and 7d**).

## MATERIALS and METHODS

### Synthesis of the chelator: synthesis of *tert*-butyl hydroxy(methyl)carbamate (A1)

N-methylhydroxylamine (2.5 g, 25.4 mmol) was dissolved in 10 mL of THF:H_2_O (1:1) and mixed with excess of K_2_CO_3_ (20.66 g, 149.5 mmol). This mixture was stirred at 0°C, using an ice bath, and a solution of Di-tert-butyl decarbonate (BOC; 7.84 mg; 35.9 mmol; THF:H_2_O 1:1) was gradually added for an hour. After the addition of BOC, the reaction mixture was left at room temperature with overnight continuous stirring. The THF was removed using Rota-evaporator and the workup of the reaction was done using Ethyl-Acetate. TLC (Hex:EtOAc/80:20) indicates the formation of the product with no starting material left. Thus, further purification was not needed. Finally, NMR validated the formation of the product. ^1^H-NMR in CDCl_3_ (400 Mhz): 1.48 (9H, s); 3.14 (3H, s).

### Synthesis of the chelator: synthesis of *tert*-butyl (benzyloxy)(methyl)carbamate (A2)

NaH (48 mg, 2 mmol) was dissolved in 2.5 mL DMF on ice, and then benzyl chloride was slowly added while waiting for the H_2_ gas to complete. After that, **A1** was dissolved in 2.5 μL DMF and progressively added while waiting for the evolution of H_2_ gas to end. Then, the temperature of the reaction was gradually raised to room temperature and stirred overnight. TLC (Hex:EtOAc /80:20) was used to monitor the reaction’s progress, and once the reaction was completed, the NaH was quenched by gently adding water. The reaction was worked up with Ethyl-Acetate and brine solution. Finally, column chromatography (Hex: EtOAc/90:10) was used to purify it. ^1^H-NMR in CDCl_3_ (400 Mhz; **Fig S1a**): 1.49 (9H, s); 3.04 (3H, s); 4.82 (2H, s), 7.39-7.34 (5H, m).

### Synthesis of the chelator: synthesis of *O*-benzyl-*N*-methylhydroxylamine (A3)

The *O*-benzyl-*N*-methylhydroxylamine (**A3**) was obtained after BOC deprotection of A2 using TFA/DCM (5:1). The reaction was fast and completed within 30 min. The TFA/DCM was evaporated by reduced pressure and diluted with diethyl ether. After that, it was washed with saturated Na_2_HCO_3_ (2 times) and brine (1 time). Finally, the pure product was obtained by drying over anhydrous Na_2_SO_4_ and concentrated under reduced pressure. ^1^H-NMR in CDCl_3_ (400 Mhz; **Fig S1b**): 2.73 (3H, s), 4.71 (2H, s), 7.38-7.34 (5H, m).

### Synthesis of the chelator: synthesis of *N*-(benzyloxy)-2-bromo-*N*-methylacetamide (Arm)

On ice, NaH (912 mg, 38 mmol) was dissolved in 10 mL of dry DCM, and then a solution of Arm (2.5 g; 18.23 mmol) was gently added. This combination was placed on ice for 30 min while waiting for the H_2_ evolution to complete. The mixture was then transferred to −78°C, and bromoacetyl bromide (4.4 g, 21.9 mmol) was added very slowly into it. The temperature of the mixture was gradually raised to room temperature, and the mixture was stirred under nitrogen. The progress of the reaction was monitored by using TLC (Hex:EtOAc /60:40) and after completion of the reaction, the NaH excess was quenched by gently adding water. With ethyl-Acetate and brine solution, the reaction was worked up. The organic layer was concentrated under reduced pressure after drying over anhydrous sodium sulfate. Lastly, the pure product was obtained by column chromatography purification (Hex:EtOAc /80:20). ^1^H-NMR in CDCl_3_ (400 Mhz; **Fig S1c**): 3.26 (3H, s), 3.90 (2H, s), 4.94 (2H, s), 7.4-7.34 (5H, m).

### Synthesis of the chelator: 1,4,7-Tris(tert-butyloxycarbonyl)-1,4,7,10-tetraazacyclododecane-10-benzyl acetate (C2a) or tritert-butyl 10-(2-ethoxy-2-oxoethyl)1,4,7,10-tetraazacyclododecane-1,4,7-tricarboxylate (C2b)

First, the 1,4,7-Tris(tert-butyloxycarbonyl)-1,4,7,10-tetraazacyclododecane (C1) was synthesized as stated in the literature[45]. Then, K_2_CO_3_ (217.4 mg, 1.575 mmol) was added to acetonitrile (10 mL) solution of tris-Boc-protected cyclen C1 (0.5 g, 1.05 mmol). The suspension was heated for 10 min at 60 °C, followed by the addition of benzyl bromoacetate (288.65 mg, 1.26 mmol) or ethyl bromoacetate (210.42, 1.26 mmol). The reaction mixture was heated for 12 h at 60–70 °C, then filtered, and the solvents removed under reduced pressure. The residue was purified by column chromatography over silica gel eluting with 10% methanol in DCM to afford the title compound as a colorless solid (5.2 g, 92% yield). ^1^H-NMR in CDCl_3_ (400 Mhz): (C2a) 7.34-7.31 (5H, m), 5.11 (2H , s), 3.62 (2H, s), 3.45-3.28 (10H, m), 2.94-2.68 (6H, m), 1.47 (27H, s); (C2b) 4.15 (2H, q, J=7.1 Hz), 3.5-3.33 (14H, m), 2.91 (4H, m), 1.45 (27H, s.)

### Synthesis of the chelator: benzyl 2-(1,4,7,10-tetraazacyclododecan-1-yl)acetate (C3) Or Ethyl 2-(1,4,7,10-tetraazacyclododecan-1-yl)acetate (C^’^3)

In this step, the deprotection of BOC was performed by using TFA. In brief, the compound C2 (500 mg) was dissolved in 200 µL of DCM. Then, 800 µL of TFA was added to the reaction mixture slowly under stirring conditions. After completion of the mixing process, the reaction mixture was left for stirring for 1.5 hours at room temperature. The TFA/DCM was evaporated by reduced pressure and diluted with diethyl ether. After that, it was washed with saturated Na_2_HCO_3_ (2 times) and brine (1 time). Finally, the pure product was obtained by drying over anhydrous Na_2_SO_4_ and concentrated under reduced pressure. ^1^H-NMR in CDCl_3_ (400 Mhz; **Fig S2a**): **C3**: 7.32 (5H, m), 3.61-3.26 (12H, m), 2.94-2.68 (6H, m).

### Synthesis of the chelator: benzyl2-(4,7,10-tris(2-((benzyloxy)(methyl)amino)-2-oxoethyl)-1,4,7,10 tetraazacyclo dodecan-1-yl)acetate (C4) or ethyl 2-(4,7,10-tris(2 ((benzyloxy)(methyl)amino)-2-oxoethyl)-1,4,7,10-tetraazacyclododecan-1-yl)acetate (C’4)

K_2_CO_3_ (10 equivalent) was added to the **C3** (500 mg, 1 equivalent) in acetonitrile (20 mL), and the reaction mixture was heated for 15 min at 65 °C. After that, the Arm (3 equivalent) was then added to the suspension, and the reaction mixture was heated at 65 °C overnight, cooled down, and the solid components were filtered and washed with cold acetonitrile. The desired product was dissolved in ultrapure water, extracted three times with dichloromethane, and the combined organic fractions were dried over Na_2_SO_4_. The organic solvent was removed by rotary evaporation to afford the product as a brown/yellow oil. The residue was then purified by column chromatography over silica gel eluting with DCM/methanol (95:5) in order to yield the title compound as a beige/white solid (60%). ^1^H-NMR in CDCl_3_ (400 Mhz; **Fig S2b**): **C4**: 7.3-7.2 (20H, m), 4.81 (8H, s), 4.18 (6H, s), 3.25 (9H, s), 3.19-1.9 (16H, m).

### Synthesis of the chelator: 2-(4,7,10-tris(2-(hydroxy(methyl)amino)-2-oxoethyl)-1,4,7,10-tetraazacyclododecan-1-yl)acetic acid (L1)

10% palladium on carbon (50 mg, 10% by weight of **C4**) was added to a solution of **C4** (500 mg, 1 eqivalent) in methanol (5 mL). After that, cyclohexadiene (10 equivalent) was added to the mixture and the mixture was heated up to 65-70° C. Finally, a balloon filled with H_2_ gas was placed on top of the mixture. The mixture was left like that for 48h. After that, the catalyst was filtered through a 0.22 μm membrane filter and washed with methanol. The solvent was removed under vacuum (reduced pressure), and the crude compound **L1** was obtained quantitatively by precipitation with diethyl ether as a white solid. ^1^H-NMR in MeOD (400 Mhz; **Fig S3a**): **L1**: 3.7-3.4 (8H, m), 3.2 (9H, s), 3.1-2.1 (16H, m). ^13^C-NMR in MeOD (150 Mhz; **Fig S3b**): 48.45 (-CH_3_), 61.83(-CH_2_), 72.39(-CH_2_), 81.63 (-CH_2_), 161.77 (-CONH), 189.63 (-COOH)

### Cell culture: human umbilical vein endothelial cell (HUVECs) culture

HUVECs were purchased from Lonza, Switzerland. HUVECs were thawed at a density of 500,000 cells into a T25 flask (Costar) and kept in culture at 37°C, 5% CO_2_ for 4 days in endothelial cell growth medium (EGM-2, Lonza). To remove DMSO from the medium while avoiding centrifugation of the recently thawed cells, cells were allowed to adhere for 4 h to the T25 flask, after which the medium was replaced. The medium was replaced every other day until a minimum of 90% confluency was achieved.

### Isolation of sEVs: hUCB-MNCs

hUCB was obtained from Daniel de Matos and Bissaya Barreto maternities, Coimbra (Portugal) (HUC-01-11). All donors signed an informed consent form according to the Portuguese legislation. Blood samples were collected into single blood bag systems containing anticoagulant citrate phosphate adenine solution. All the isolation process was following the previously published protocol [17]. Briefly, One to 2-day-old blood was diluted 1:1 with a dilution buffer (1× PBS (GibcoTM, Thermofisher Scientific) with 2 mM ethylenediaminetetraacetic acid (EDTA, Alfa Aesar)). This diluted blood (30-35 mL) was then layered over 15 mL Lymphoprep^TM^ (STEMCELL Technologies Inc.) and centrifuged at 400 g for 35 min (20°C, without brakes) to separate the different components of the blood. The mononuclear cell layer, called buffy coat, characterized by a white cloudy layer of cells that stands between the layer of lymphoprep (bottom) and the layer of plasma (top), was carefully removed and transferred to new falcon tubes, which were centrifuged at 300 g for 15 min (RT). The resulting supernatant was discarded, and the cell pellet was resuspended in MACS buffer (dilution buffer with addition of 0.5% Bovine Serum Albumin (BSA, Sigma-Aldrich)) and centrifuged at 300 g for 10 min (RT). The supernatant was once again discarded, and the cell pellet was resuspended in 50 mL of MACS Buffer. Ten µL were removed for cell counting, and the rest of the volume was centrifuged at 200g for 12 min (RT). The cell pellet was resuspended in freezing medium (70% cells in AIM V medium (Gibco), 20% FBS, and 10% DMSO) at a density of 100 M cells per vial and stored at −80°C within a gradual freezing container.

For EV isolation from MNCs, cells (2 million cells per mL) were seeded into each well of a 6-well plate (Costar) in AIM V^TM^ (Gibco) medium supplemented with stem cell factor (50 ng/mL, SCF, Petrotech), fms like tyrosine kinase 3 (50 ng/mL, FLT-3, Petrotech) and DNAse I (50 μg/mL, Sigma Aldrich Co) and incubated in a hypoxia chamber (BioSpherix culture chamber, BCA Scientific) under humidified atmosphere with 0.5% O_2_ and 5% CO_2_, at 37°C. After 18 h, the cell culture medium was collected and EVs isolated by differential ultracentrifugation.[46] First, conditioned medium was centrifuged at 300 g for 10 min at 4°C to pellet cells, and supernatant was again centrifuged at 2000 g for 20 min, also at 4°C to pellet cell debris. Using an Optima™ XPN 100K ultracentrifuge with a swinging bucket rotor SW 32 Ti (Beckham Coulter), the supernatant was two times centrifuged at 10,000 g for 30 min at 4 °C in order to pellet microvesicles. EVs were then pelleted with a 100,000 g centrifugation for 120 min at 4°C. Pelleted EVs were washed with cold PBS, recentrifuged at the same speed, resuspended in 150-200 μL of cold sterile PBS and preserved at −80°C.

### Isolation of sEVs: urine

Urine samples were obtained from healthy donors upon signing an informed consent (ref: CE-070/2019; Faculty of Medicine, University of Coimbra). EVs were isolated from urine samples as previously described [47]. Briefly, the urine sample was vortexed for 90 sec, centrifuged for 20 min at 2000g, and then the supernatant was diluted in Tris-EDTA (4 times, 20 mM, pH 9). The centricon® Plus-70 centrifugal filter devices were sanitized with 70% EtOH and centrifuged at 3500 g for 10 min. The diluted urine samples were filtered using a 0.22-µm pore-sized filter with a PES membrane. Then, 50 mL was added to each centricon® Plus-70 centrifugal filter device and centrifuged for 12 min at 25°C, and repeated until all the sample was processed. Next, the top part of the centricon filters was attached to the collection cups and centrifuged at 1000g for 2 min. Further, this EV preparation was purified using qEV size exclusion chromatography (SEC) columns and finally concentrated using an ultracentrifuge (100 000g, 2 h, 4 °C).

### Labelling of sEVs: CFSE labelling

The lumen of EVs was labelled with the CellTrace™ CFSE Cell Proliferation Kit (ThermoFisher Scientific). CellTrace™ CFSE in DMSO (5 mM) was added to the EV sample to a concentration of 20 µM and incubated for 90 min at 37°C with agitation. After the incubation, 1% BSA/PBS that was previously filtered (0.2 µm Polyethersulfone membrane filter) was used to block the solution for 10 min (RT, at a concentration of 10% of the EV solution volume). Following blockage of the solution, EVs were purified with qEV SEC and then concentrated by ultracentrifugation for 2 h at 100,000g (4°C) in ultracentrifuge tubes. Finally, EVs were resuspended in 110 µL of PBS and stored at −80°C.

### Labelling of sEVs: Zr^4+^ labelling

For labelling the EVs with L1, EDC/NHS chemistry was used. Briefly, the -COOH group in our chelator L1 was activated using 10 times excess EDC (1-ethyl-3-(3-dimethylaminopropyl)carbodiimide) followed by the esterification using sulfo-NHS (2.5-time excess) at pH 5.5 using MES buffer. After 15 min, the pH of the mixture was increased to pH 7.5, mixed with our EVs, and incubated at 37°C for 2 h. The excess chelator was removed by using the Exo-spin column. Finally, the presence of L1 on the EV surface was confirmed by the reduction of the number of amine groups after L1 modification. Then, 2.5×10^10^ −3.0×10^10^ EV-L1 was incubated with ^89^ZrCl_4_ (∼1.5 mCi), which was obtained from ICNAS [48, 49] at 37° C for 1 h. The ^89^Zr-oxalate was supplied by ICNAS, which was converted to the ^89^ZrCl_4_ by using Waters Sep-pak Light accell plus QMA strong anion exchange cartridge using standard protocol [50]. After that, the excess volume of the ^89^ZrCl_4_ was removed by evaporation, and the pH was adjusted close to ∼6.0. Finally, the solution of the EV-L1 was added to the ^89^ZrCl_4_ in tris buffer with pH ∼ 7.4 and incubated for 1 h at 37°C. The ^89^Zr^4+^ excess was removed by the Exo-spin column, and the purity was measured by iTLC. **Characterization of sEVs:** nanoparticle tracking analysis (NTA). EV size and concentration analysis was performed using the NanoSight NS300 (Malvern Instruments, Malvern, UK). EVs were diluted to a desired concentration of between 20 and 40 particles per frame in 1x PBS. 1 mL of this diluted sample was loaded into the equipment using a syringe, and then 5 videos of 30s each were acquired, using a camera level of 14-15. The resulting videos were processed and analysed using NTA 3.4 analytical software, with threshold values chosen according to the quality of the video and of the sample, which stood between 4 and 6.

### Characterization of sEVs: microBCA^TM^ analyses

Micro BCA^TM^ protein assay kit (Thermo Fisher Scientific) was used to quantify the total protein amount in EVs. EVs were diluted in 2% sodium dodecyl sulphate (SDS) in order to disrupt EVs membrane. The protein quantification was done against a standard BSA curve. Dilutions were prepared following manufacturer’s instructions. Absorbance values were read in a microplate reader at 562 nm after 2 h incubation at 37° C, followed by a 15 min incubation at RT.

### Characterization of sEVs: zeta potential analyses

Zeta potential of EVs was measured (Zeta PALS potential analyzer, Brookhaven Instruments Corporation) by diluting 4 µL of EV stock in 1.5 mL of MiliQ Water (4.0 ×10^9^ – 1.2 × 10^10^). The conductance of the solution was adjusted to 250-300 µS by using a 3M KCL solution. Zeta potential measurement was conducted for 5 runs.

### Characterization of sEVs: transmission electron microscopy (TEM)

EV samples (5 µL) were fixed in 2% paraformaldehyde (PFA) (w/v) and then layered over EM-Tec formvar-carbon support film on copper 300 square mesh grills (Micro to Nano Innovative Microscopy Supplies) for 5 min at RT. These grids were then turned upside down on a drop of uranyl-acetate 2%, a contrasting agent, for 1 min at RT. TEM images were taken using a Tecnai G2 Spirit BioTwin electron microscope (FEI).

### Characterization of sEVs: quantification of amine groups on sEV surface

The number of amines on the surface of EVs was quantified by using the fluorescamine assay[51, 52]. Fluorescamine is a non-fluorescent spiro compound that reacts with primary amines to form highly fluorescent products. Unreacted Fluorescamine reagent and its degradation products are non-fluorescent.

### *In vitro* biological assays: cytotoxicity of EV-L1-Zr

HUVECs in P8 were plated at a cell density of 8,000 cells per well in a MW96 Corning^®^ Costar^®^ cell culture plate (Corning Inc.) and kept in culture for 20 h. The cells were then treated with EVs (1×10^9^ part/mL native MNC SEVs), Zr^4+^-labeled EVs (1×10^9^ part/mL MNC EV-L1-Zr and EV-DFO-Zr), and Zr^4+^ (10 nM) in EGM-2 containing 2% FBS (EV-depleted). The medium from the non-treated cells control was replaced with EBM-2, while the medium from the positive control condition was replaced with EGM-2 medium. The plate was then incubated for 48 h. At the end of this timepoint, cells were incubated with Hoechst (1 µg/mL in EGM-2) for 30 min at 37°C and then imaged using a high-content fluorescence microscope (IN Cell 2200, GE Healthcare). Finally, the images were analyzed automatically in IN Cell Developer software (GE Healthcare) by nuclei count.

### *In vitro* biological assays: cellular internalization of EV-L1-Zr

HUVECs in P8 were plated at a density of 6,000/well, in a µ-Slide 15 Well (IBIDI) plate and cultured in EGM-2 media for 24 h. After this, both CFSE-labelled sEVs, sEV-L1-Zr and sEV-DFO-Zr, were added to the plated cells at a concentration of 1×10^11^ particles/mL in EGM-2 + 2% FBS (EV-depleted). Control condition cells just had their medium replaced with the same medium as the other conditions. The cells were then incubated for 4 h to promote the internalization of sEVs or EV-L1-Zr or EV-DFO-Zr, after which the cells were incubated with LysoTracker^TM^ Red DND-99 (Invitrogen, 100nM) and with Hoechst (0.5 µg/mL) for 30 min. After this incubation, cells were washed with 1x PBS to remove non-internalized SEVs and were then fixed with 2% PFA for 10 min and then kept in 1x PBS at 4°C overnight. The next day, cells were imaged with a Zeiss LSM 710 confocal microscope (Carl Zeiss), using the lasers 488, 405, and 561, with a Plan-Apochromat 40×/1.4 oil immersion objective.

For cellular internalization studies by flow cytometry analyses, HUVECS in P8 were plated onto a MW24 cell culture plate (Corning Inc.) at a density of 60,000 cells per well. The cells were cultured for 20 h and then treated with CFSE-EVs (1×10^11^ part/mL urine CFSE-EVs) or CFSE-EV-DFO-Zr (1×10^11^ part/mL urine CFSE-EV-DFO-Zr) and Zr (10 nM) in EGM-2 containing 2% FBS (EV-depleted) for 4 h. Control cells had the medium replaced with EGM-2, including 2% FBS (EV-depleted). After the 4 h period, the medium was removed in all the conditions, and the cells were incubated for 10 min at room temperature with Trypan Blue/PBS (500 µL at 0.04%) to promote quenching of the fluorescence signal outside the cells. The cells were then trypsinized (0,5% Trypsin-EDTA), centrifuged at 300 g for 5 min, and finally resuspended in 1xPBS (200 µL) and kept on ice until acquisition. The flow cytometry analyses were performed in a BD Accuri C6 Flow Cytometer (BD Biosciences) and the data were analyzed with the FlowJo^TM^ (v10, FlowJo, LLC) software. A minimum of 10,000 events was collected, and the resultant percentages of the analysis were calculated based on the negative cell control (gate was defined for this control at 1%).

### *In vitro* biological assays: cell survival properties of EV-L1-Zr

The bioactivity of the Zr^4+^ labeled EVs was evaluated by using a survival assay [53, 54]. Briefly, HUVECs in P8 were plated at 8000 cells/well in a MW96 Corning^®^ Costar^®^ cell culture plate (Corning Inc.) in EGM-2 medium (LONZA) and kept at 37°C (5% CO2) for 20 h. Then, the medium was removed and replaced with EGM-2 (LONZA) containing 2% FBS (EV-depleted), and the cells were treated according to the following description (i) control (EGM-2 (LONZA) containing 2% FBS (EV-depleted); (ii) positive control (EGM-2 instead of EGM-2 (LONZA) containing 2% FBS (EV-depleted); (iii) EVs (1×10^9^ part/mL native MNC SEVs); (iv) Zr^4+^ labeled EVs (1×10^9^ part/mL MNC EV-L1-Zr, EV-DFO-Zr); (v) Zr (10 nM). The plate was then incubated at 37°C (5% CO_2_) for 48 h. At the end of this period, the medium was removed and replaced with endothelial basal medium (EBM-2, Lonza) containing gentamycin and amphotericin B, in all conditions except for the positive control condition, in which the medium was refreshed with EGM-2 medium. Then the cells were incubated in a hypoxic environment (0.1% CO_2_, 37°C) for another 48 h. After this, the medium was removed, and we incubated the cells with Hoechst 33342 (Invitrogen, 1 µg/mL in EGM-2) for 30 min (nuclei staining). Images were then acquired using a high-content fluorescence microscope (IN Cell 2200, GE Healthcare) and analyzed automatically in IN Cell Developer software (GE Healthcare) by nuclei count.

### *Ex-vivo* Langendorff heart perfusion model (LHP): setup and testing

LHP was produced from the hearts collected from 10-week-old Wistar Han rats (400±25 g). The animals were anaesthetized with 85 mg/kg ketamine and 10 mg/kg xylazine and heparinized. Hearts were perfused on a Langendorff apparatus [perfusion pressure of 9333 Pa (70 mmHg), constant flow rate of 2.5×107 m^3^ /s (15 mL/min)], with modified Krebs–Henseleit (KH) buffer (118 mmol/L NaCl, 25 mmol/L NaHCO_3_, 4.7 mmol/L KCl, 1.2 mmol/L MgSO_4_, 1.2 mmol/L KH_2_PO_4_, 10 mmol/L HEPES, 1.25 mmol/L CaCl_2_ and 10 mmol/L glucose, pH 7.4), equilibrated with 95% O_2_/5% CO_2_ at 37°C. Perfusion was let to stabilized for 10 min, after which hearts were either perfused for a further 60 min mixed with PBS (controls), or 7.5 × 10^9^ EVs labelled with Zr^4+^ or DiL (fluorescence dye, Invitrogen^TM^). After the incubation with sEVs, a 10 min washing step was performed to remove the excess of EVs from the heart.

### LHP: ICP-MS analysis

For isolation of rat cardiomyocytes/non-cardiomyocytes, after the LHP was done following our previously optimize method [35]. Briefly, after the indicated treatments, hearts were perfused with digestion buffer (130 mM NaCl, 5.4 mM KCl, 3 mM Pyruvate, 25 mM HEPES, 0.5 mM MgCl_2_, 0.33 mM NaH_2_PO_4_, 22 mM Glucose, 10 mM BDM and 1 mM EGTA) for 2 min. Subsequently, hearts were perfused with collagenase digestion buffer (2.4 mg/ml type II collagenase (Gibco), 0.06 mM CaCl2) for 20 min, followed by mechanical digestion for 3 min, using forceps and plastic Pasteur pipettes in stopping solution (1.2 mg/ml type II collagenase (Gibco), 0.12 mM CaCl_2_, 10 mg/mL BSA). Cell suspension was filtered through a 300 μm mesh filter and centrifuged at 100 rpm, for 1 min. Viable cardiomyocytes were isolated following resuspension in CaCl_2_ buffers of increasing Ca^2+^ concentrations to reach a final concentration of 1 mm Ca^2+^. Supernatants from all centrifugations were pooled, filtered through a 70 μm cell strainer and centrifuged at 1000 g for 5 min, to isolate non-cardiomyocytes. Typically, from a rat heart we isolated approx. 1-10 ×10^6^ cardiomyocytes and 3-6.5×10^6^ non-cardiomyocytes. Subsequently, both CM and non-CM cells were frozen in liquid nitrogen, followed by freeze-drying. Samples were digested with 2 ml of HNO_3_ 69% (Hiperpur, Panreac) and 1 mL H_2_O (MilliQ) in a microwave system (Milestone Ultrawave) at 240°C for 20 min. After digestion, MilliQ water was added until 25 mL final volume. The resulting samples were digested with HNO₃ and analysed using ICP-MS (Agilent 7700x) equipped with an introduction system composed of a Micromist glass low-flow nebulizer, Scott spray chamber with Peltier (2°C), and a quartz torch. For the percentage calculation, the amount of Zr^4+^ in the Krebs buffer was considered as the total Zr^4+^.

### LHP: heart sections

Briefly, after the LHP the rat hearts were cut into half in transverse position and frozen in OCT solution and stored at −80° C. Hearts were mounted using OCT cryomatrix (Thermo fisher scientific, USA) and cryosections of 15 µm in the coronal plane were obtained using a Leica CM1950 cryostat and collected to glass slides (Superfrost plus, VWR, USA), one representing each region per slide. Slides were kept at −80°C until labelling. **LHP: immunohistochemistry.** Frozen glass slides containing heart sections were thawed at room temperature and rehydrated by soaking in PBS for 10 min. Heart sections were contoured using a hydrophobic pen (Immedge™ Hydrophobic Barrier Pen, Vector Laboratories, ImmEdge™), and the sections were fixed using 4% PFA. After that the sections were incubated with 0.3% of triton for 15 min at room temperature followed by blocking with 3% BSA/ PBS and incubation with monoclonal anti-α-actinin antibody (A7811, Sigma-Aldrich) (1: 200) diluted in 1% BSA/ PBS for overnight at 4°C. Then, the slices were washed 3 times with PBS and incubated with Alexa Fluor 633 goat anti-mouse (1: 500) (ThermoFisher) for 2 h at room temperature. Finally, the slice was washed 3 times with PBS followed by staining with Hoechst 33342 (2 µg/ mL, Invitrogen). The slice was then embedded using Mowiol 40-88/ glycerol mounting medium. Three z-stack N(8 “Z”) images were acquired per cardiac slice on a Zeiss LSM 710 confocal microscope (Carl Zeiss) using a 100× objective/ 1.4 numerical aperture oil Plan Apochromat immersion lens. sEVs were identified and counted by the increased gray levels compared to the background using the Analyse Particles plugin on ImageJ software. Identification of the corresponding cell was then manually assessed by colocalization with cardiac α-actinin staining (cardiomyocyte). The cells without the α-actinin staining were considered as a non-cardiomyocytes and the sEVs close to those cells’ nuclei were considered as non-cardiomyocytes accumulated EVs. The distribution of sEVs within the heart slices was quantified by capturing five images per slice. Z-stack images (comprising eight sections) were acquired using a Zeiss LSM 710 confocal microscope (Carl Zeiss) equipped with a 40× oil immersion objective lens with a 1.4 numerical aperture (Plan Apochromat). Quantitative analysis was performed utilizing QuPath and ImageJ software.

### *In vivo* studies: biodistribution and PET/MRI imaging

All PET scans were performed using a prototype of a high-acceptance small-animal PET based on resistive plate chambers (RPC-PET) [55]. All mice underwent the PET acquisition with the radiopharmaceuticals sEV-L1-Zr and sEV-DFO-Zr. The PET acquisition lasted for 24 h post-injection. Images were reconstructed using OSEM algorithm and a cubic voxel of 0.5 mm width.

Five fiducial markers that can be viewed both in the PET and MRI imaging were placed in the mouse bed, and MRI imaging was done after PET without moving the mice from the bed, under anesthesia. Mice were kept anesthetized by isoflurane (1.5-2.0%) with 100% O_2_ , with body temperature and respiration monitoring (SA Instruments SA, Stony Brook, USA). All MR experiments were performed in a BioSpec 9.4T MRI scanner (Bruker Biospin, Ettlingen, Germany) with a volume resonator (transmitter/receiver). Morphological images were acquired with localizer-multi-slice in axial, saggital and coronal orientations, and a 2D T2-weighted turbo RARE sequence in axial orientation. The localizer sequence had the following parameters: TE/TR=1.901/288 ms, FOV=45.0*45.0 mm, acquisition matrix=226*226, averages=5, flip angle=30.0°, 70 axial continuous slices with 1.0 mm thick and acquisition time of 2m44s, with respiratory gating. The 2D T2-weighted turbo RARE sequence had the following parameters: TE/TR=11/3100 ms, FOV=70.0*45.0 mm, acquisition matrix=350*220, averages=1, rare factor=1, echo spacing=11 ms, 34 coronal continuous slices with 0.8 mm thick and acquisition time of 11m22s.

### *In vivo* studies: Zr uptake quantification

First, based on the fiducial marker, the images obtained from the PET were manually registered by the MRI images. For quantitative PET imaging processing, PMOD and FUSION Tool (PMOD 3.6; PMOD Technologies; Zürich, Switzerland; RRID:SCR_016547) were used to delineate volumes of interest (VOIs) on the PET images co registered with the respective MRI. Regions of interest (ROIs) were defined manually based on the anatomical MRI images. Uptake values are expressed as percentage of injected dose per gram of tissue (% ID/g).

## Author Contributions

Design and synthesis of the chelator: AB; ^89^Zr production and radiolabelling of the sEVs: AB, IH and MS; PET Imaging and Bio-distribution: JS, IH, and AB; In-vitro assays and microscopic images: AB and CJ; LHP model: AB, TM and HG; Data interpretation: AB, IH, JS, TM, HG, MJF, AA and LF; Writing the manuscript: AB, LF, AA; Supervision of the project: LF.

## Acknowledgments

This work was supported by Project ‘LABEL’ (POCI-01-0247-FEDER-049268) under the Portugal 2020 program, FCT project reference TargetingHeart 2022.02803.PTDC and 2023.15026.PEX, Pizo4spine- grant agreement No 101098597 (European Union’s Horizon Europe research and innovation programme), Reborn- 101091852 (European Union or the European Health and Digital Executive Agency (HADEA)). Carlos Jesus is thankful for his FCT fellowship (SFRH/BD/144092/2019).

## GEO Location

This study was conducted at the University of Coimbra, Portugal.

---

**Scheme 1.**
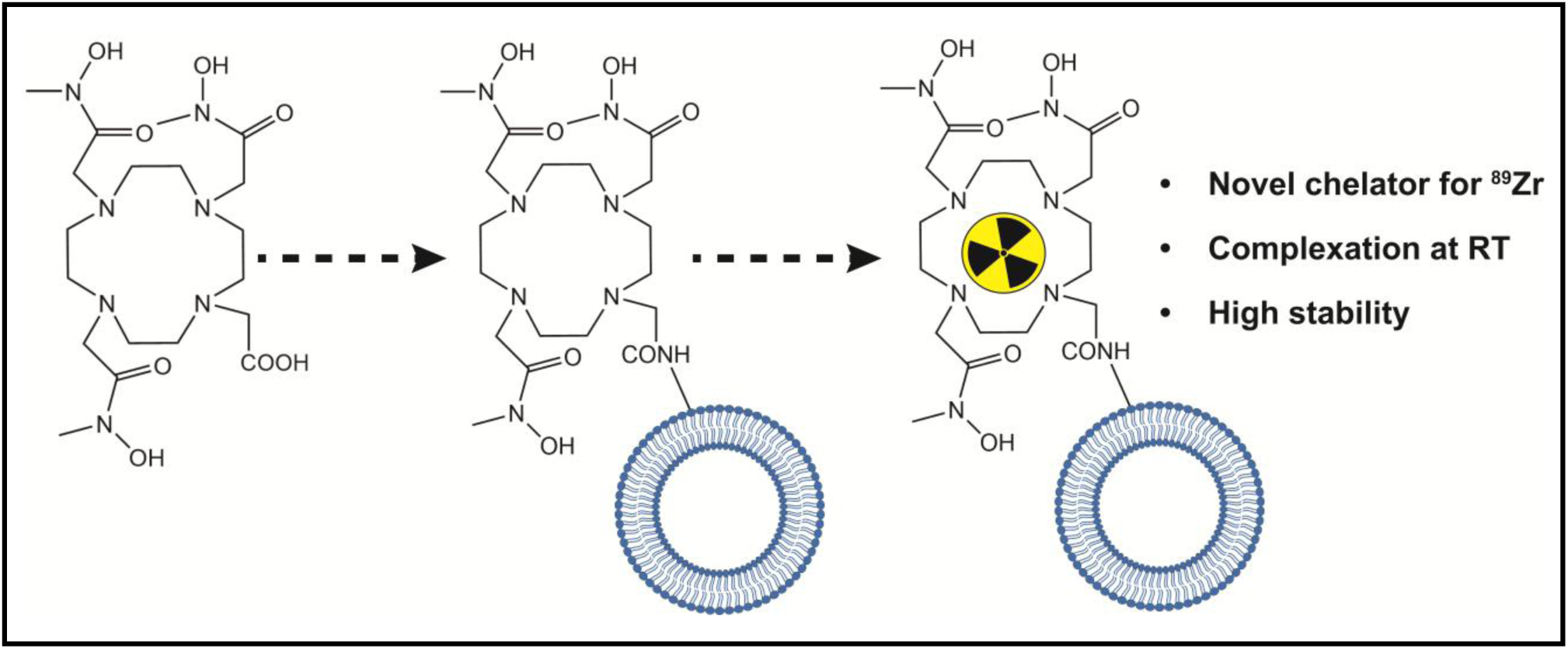
Schematic presentation of the present work. This work describes the design of a novel chelator of Zr^4+^ that forms a complex at room temperature and provides higher stability than conventional chelators.

