## Supplemental figure and table for "A room-temperature ⁸⁹Zr⁴⁺ radiolabelling strategy for small extracellular vesicles with enhanced plasma stability for PET Imaging"

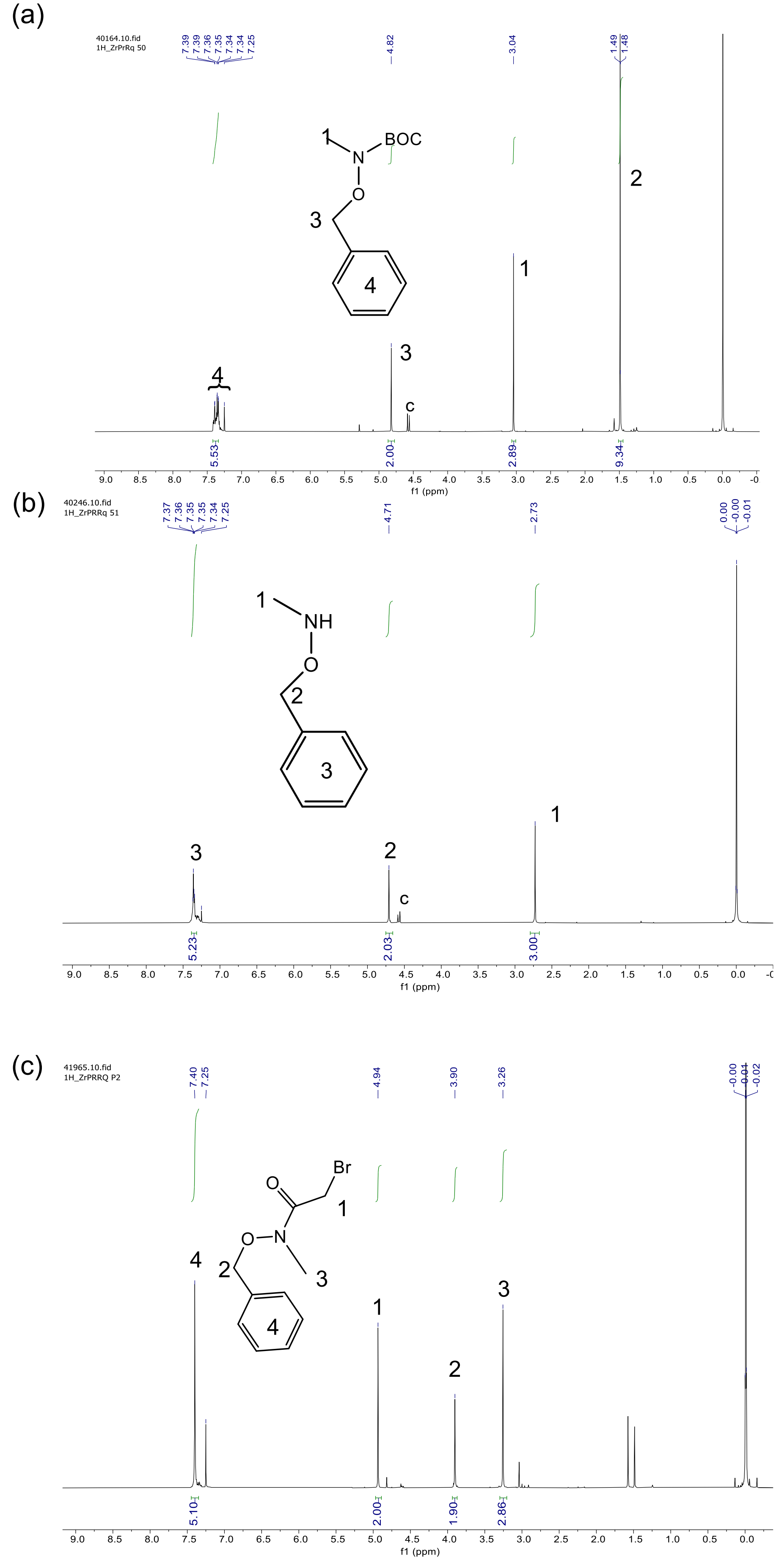


**Figure S1.** $^{1}$H-NMR of *tert*-butyl (benzyloxy)(methyl)carbamate (A2), *O*-benzyl-*N*-methylhydroxylamine (A3) and *N*-(benzyloxy)-2-bromo-*N*-methylacetamide (Arm).

(a)

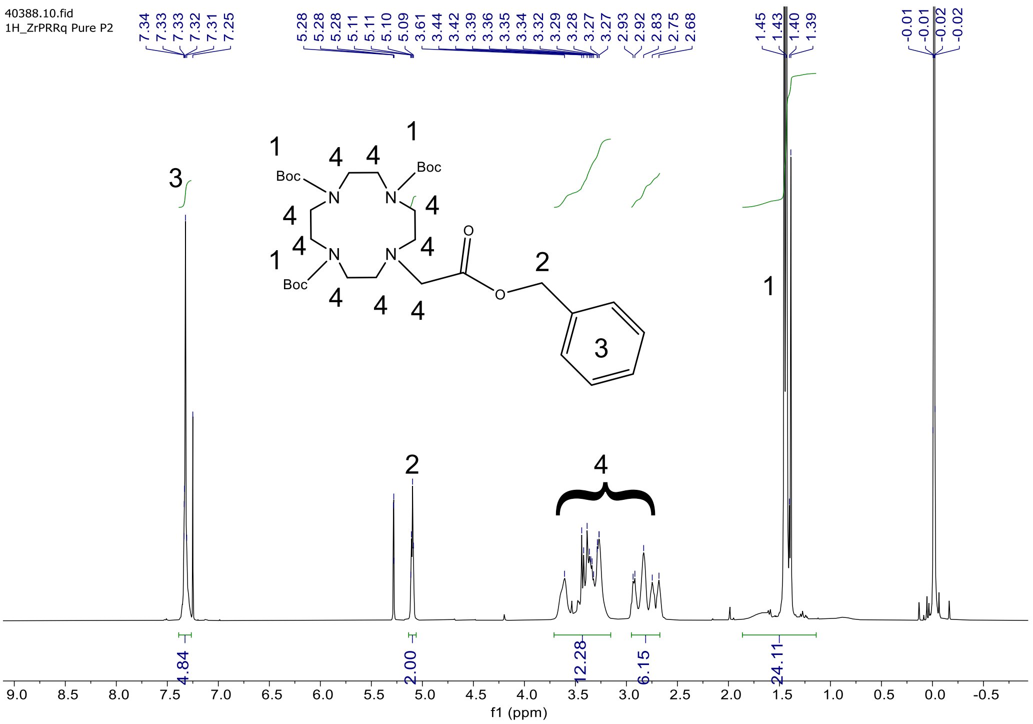


(b)

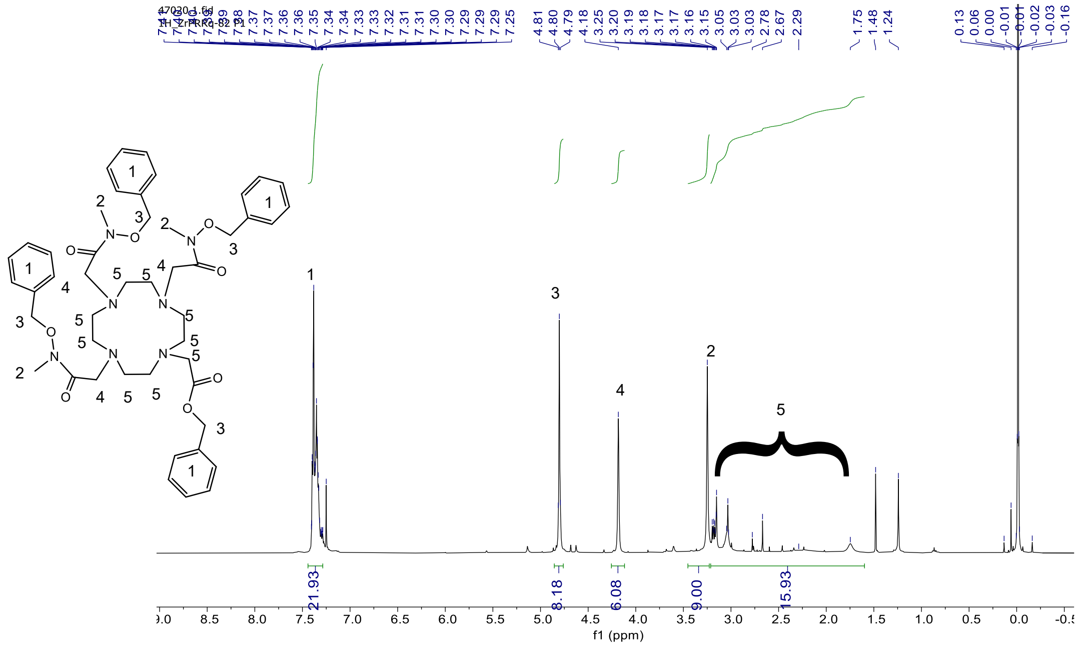


**Figure S2.** [1]H-NMR of 1,4,7-tris(tert-butyloxycarbonyl)-1,4,7,10-tetraazacyclododecane-10-benzyl acetate (C2) and benzyl 2-(1,4,7,10-tetraazacyclododecan-1-yl)acetate (C3).

(a)

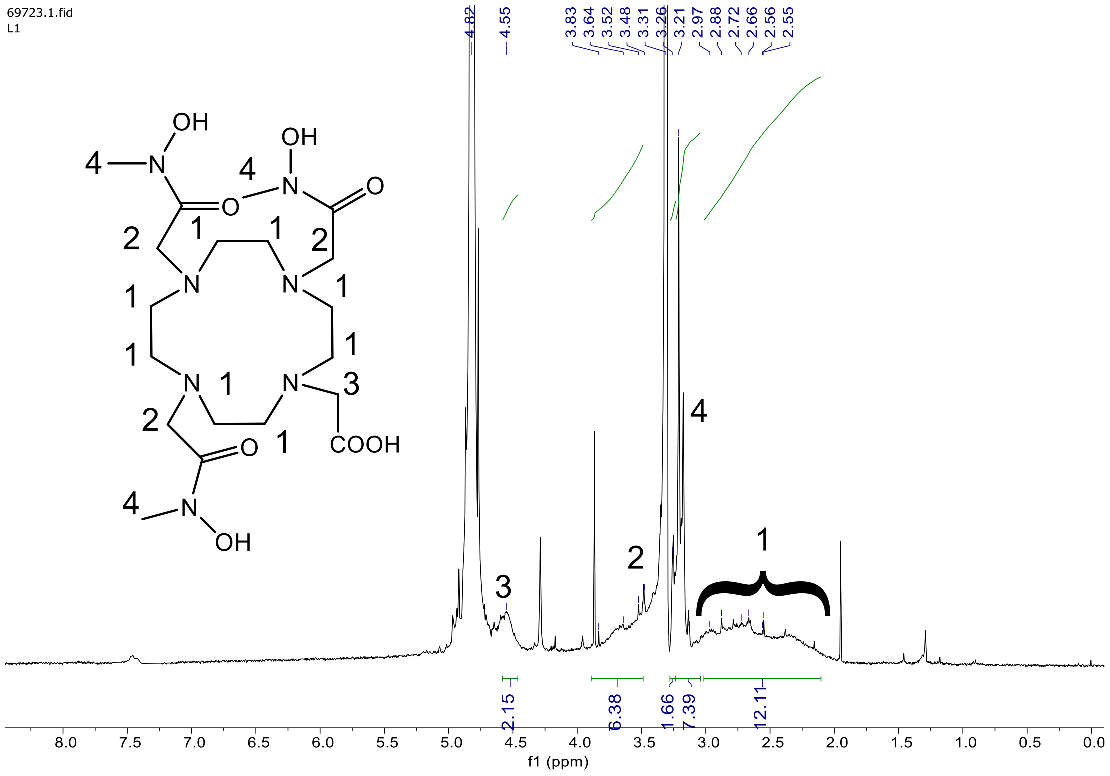


(b)

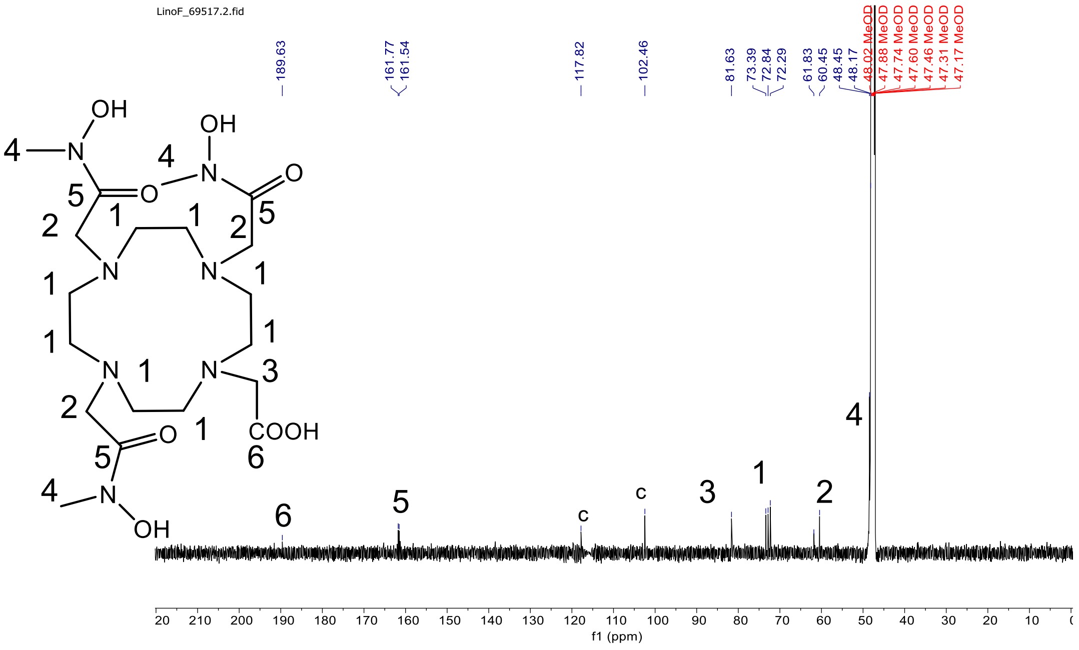


**Figure S3**. $^{1}H$-NMR of 2-(4,7,10-tris(2-(hydroxy(methyl)amino)-2-oxoethyl)-1,4,7,10-tetraazacyclododecan-1-yl)acetic acid (L1) and $^{13}C$-NMR of 2-(4,7,10-tris(2-(hydroxy(methyl)amino)-2-oxoethyl)-1,4,7,10-tetraazacyclododecan-1-yl)acetic acid (L1).

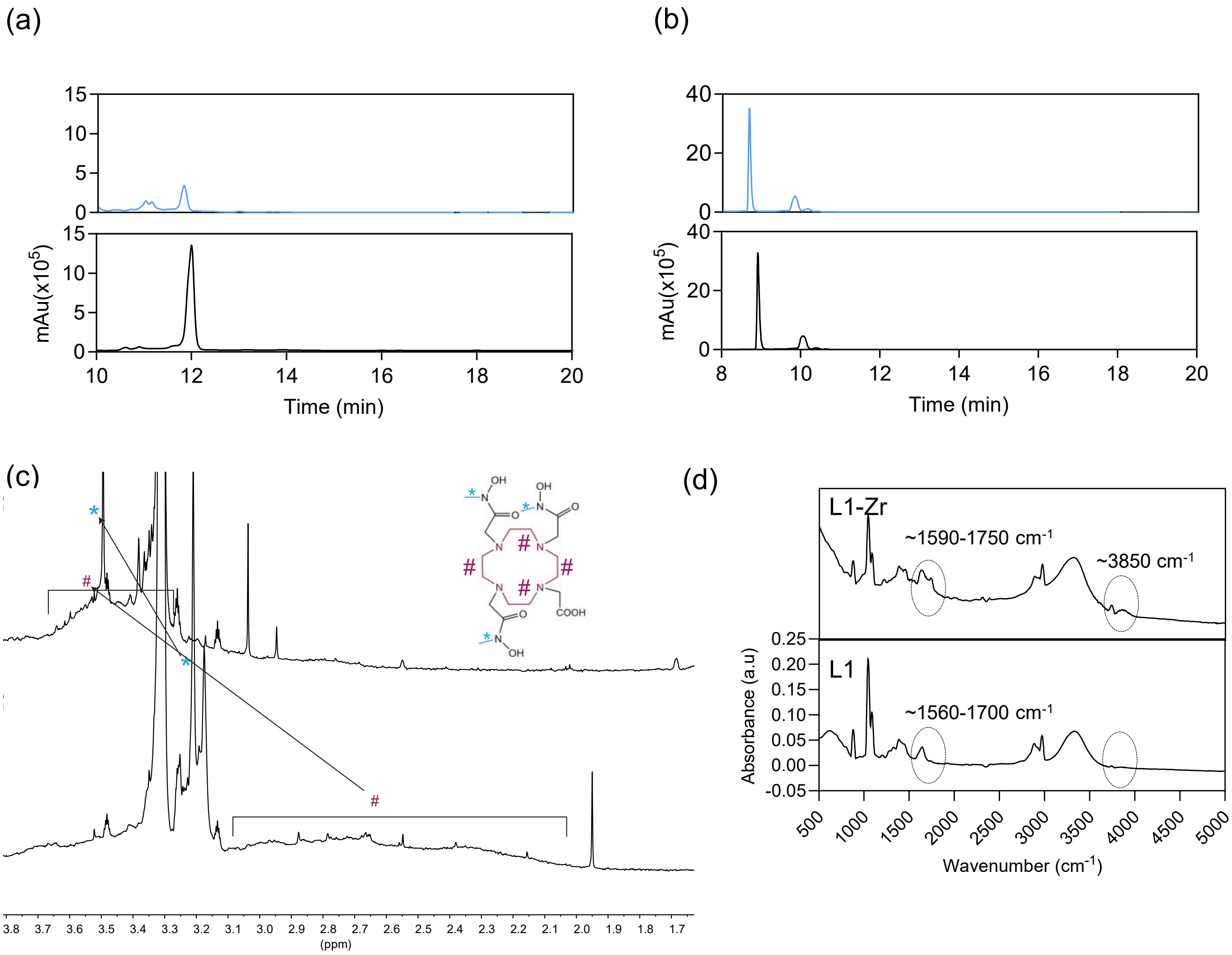


**Figure S4. Complexation of $Zr^{4+}$ with various chelators (DFO, DOTA and L1).** (a) HPLC chromatogram of DFO and DFO-Zr. (b) HPLC chromatogram of DOTA and DOTA-Zr. (c) Chemical shift in the $^{1}H$ due to the formation of the L1-Zr complex. (d) FTIR shift due to the formation of the L1-Zr complex.

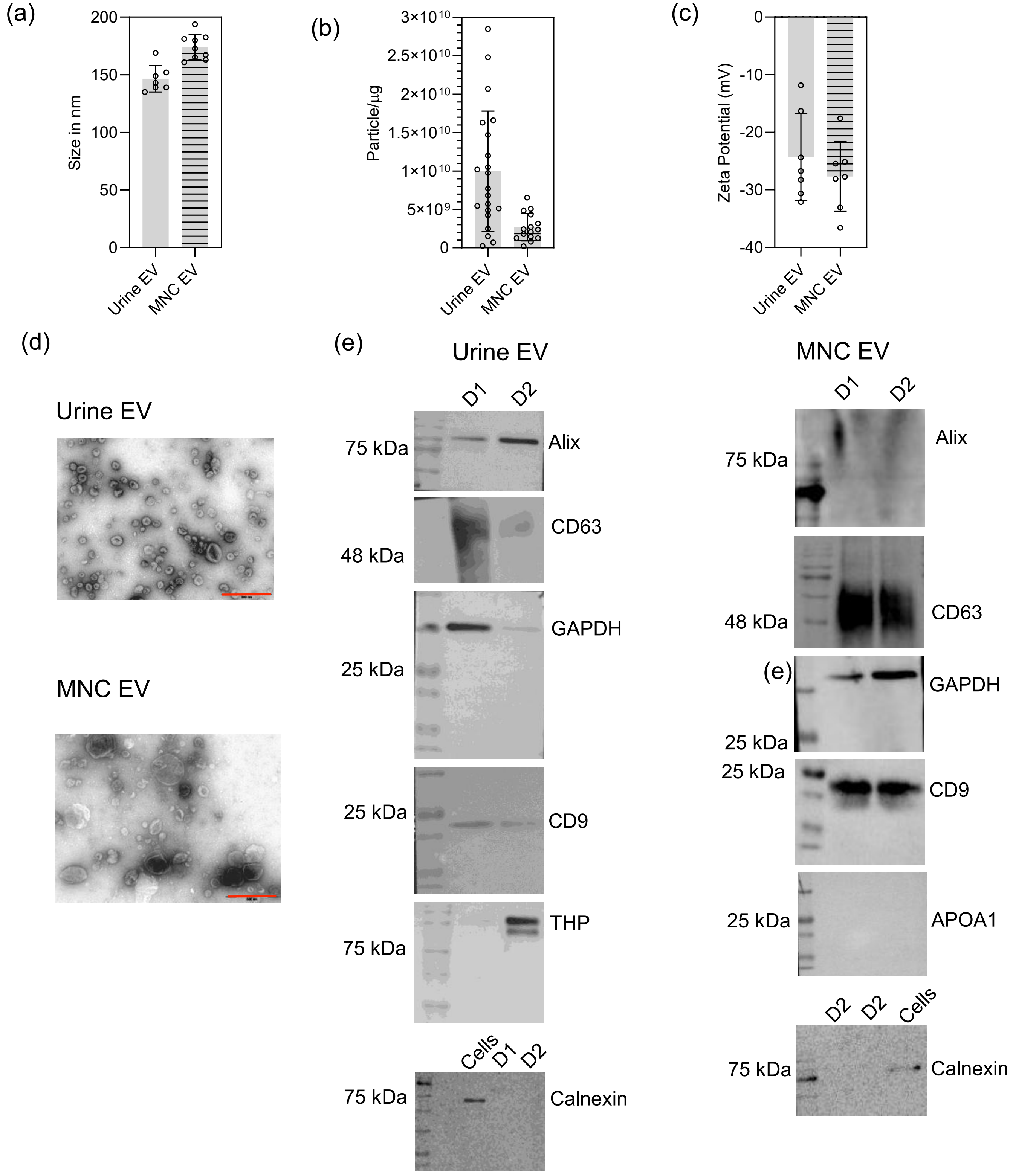


**Figure S5. Characterization of sEVs isolated from different sources.** (A) Mean size of sEVs isolated from urine and MNCs. (B) Particle per µg of protein of sEVs isolated from urine and MNCs. (C) Zeta potential of the sEVs isolated from urine and MNCs. (D) TEM images of sEVs from urine and MNCs. The red bar indicates 500 nm. (E) Western blot analyses of sEVs from urine and MNCs.

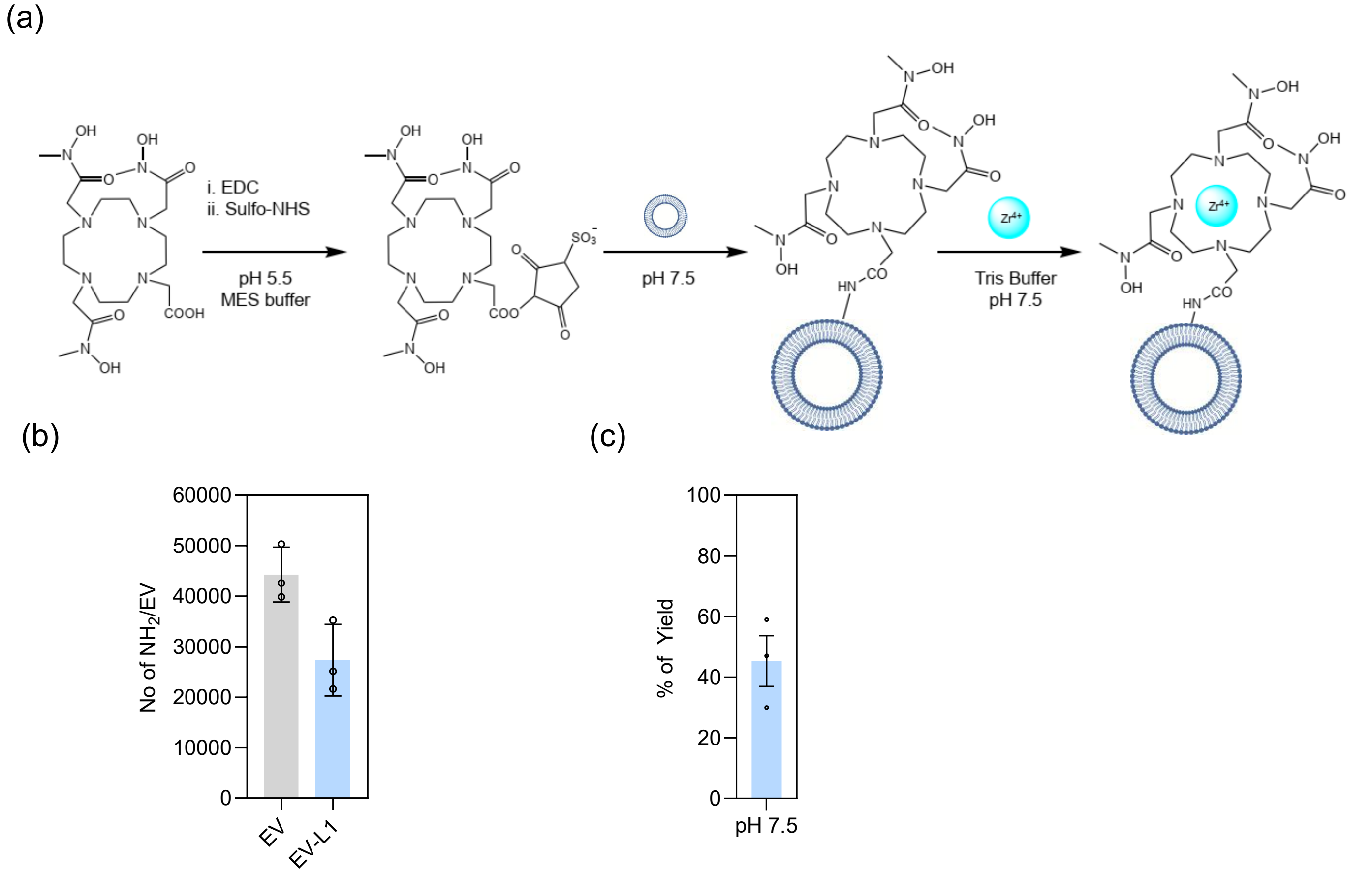


**Figure S6. Labelling sEVs with $Zr^{4+}$using L1 as a chelator.** (A) Schematic illustration of sEV labeling with $Zr^{4+}$. This process occurs in two steps: first, sEVs are labeled with the chelator L1, followed by the addition of $Zr^{4+}$ for the subsequent labeling. (B) Number of amine group quantifying before and after the labelling with **L1** using a fluorescamine assay. (C) Yield of sEV labeling at pH 7.5.

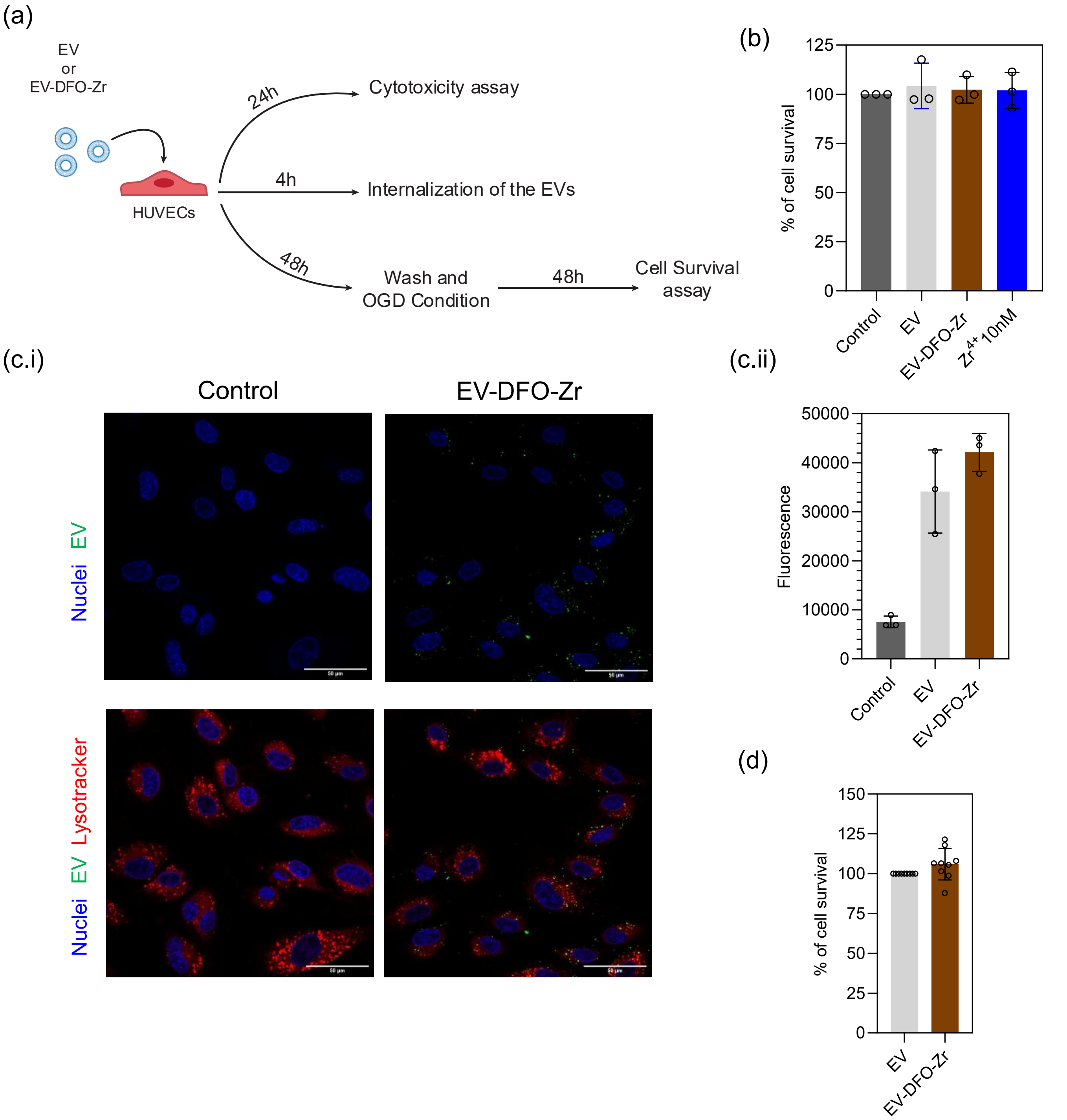


**Figure S7. Cytotoxicity, cellular internalization and bioactivity of EV-DFO-Zr.** (a) Schematic representation of the protocol used to assess internalization of sEVs and sEV-DFO-Zr using confocal and flow cytometry. Colocalization of CFSE-labelled sEVs and sEV-DFO-Zr with Lysotracker was imaged using confocal microscopy. (b) Cytotoxicity measurements of sEV, sEV-DFO-Zr and Zr (10 nM) on HUVECs. (c) Internalization of sEVs and sEV-DFO-Zr on HUVECs. Bars represent mean ± SD. Scale bars: 33 µm. (d) Bioactivity of sEV-L1-Zr (see Materials and Methods). % of endothelial cell survival when treated with sEV, sEV-DFO-Zr, $Zr^{4+}$ or EGM (containing VEGF) and then exposed to hypoxia. In b-d, results are Mean ± SD (n=3 independent runs and 3 technical replicates per independent run). Asterisks (**; ***) denote statistically significant difference (Ordinary one-way ANOVA with Dunnett's multiple comparisons against control; p< 0.01; p< 0.001, respectively).

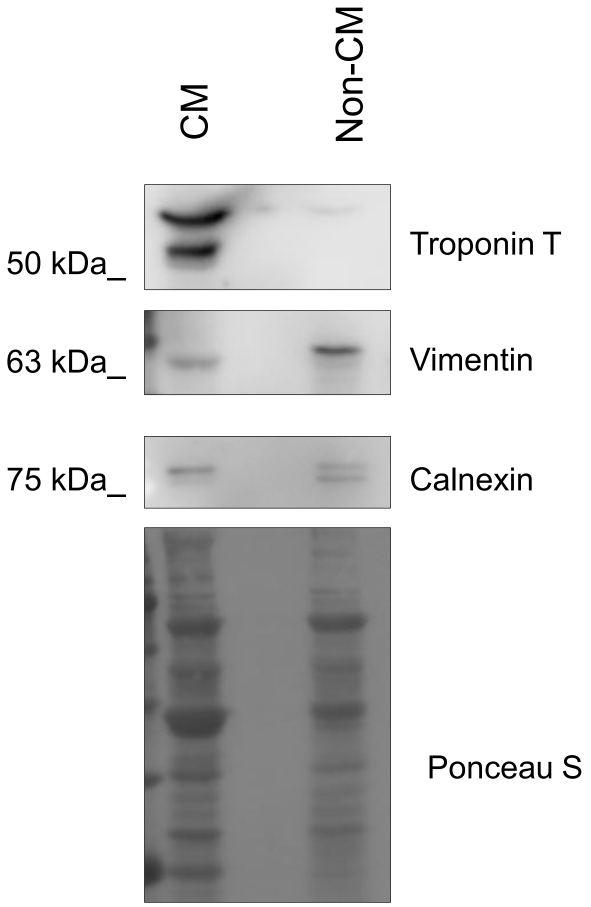


**CM: cardiomyocytes**

**non-CM: non-cardiomyocytes**

**Figure S8. Characterization of cardiac cells isolated from the Langendorff model.** Characterization by Western blot of CM and the non-CM cells isolated from the Langendorff model. Troponin T was used as a marker for the CM.

**Supplemental Table 1. Comparison of sEV-L1-Zr with previous work**

| | pH for labeling | Yield | Purity | Stability | Maximum number of Zr per sEV | Bone accumulation |
|---|---|---|---|---|---|---|
| Zr-Oxinate intraluminal labelling | 7.4 | 6.2% (MDA-MB-231 sEV) 16.2% (PANC1 sEV) | Did not report | In PBS at 37ºC 75% (24 h), ~ 40% (48 h) | Not Reported | ~10%: |
| sEV-DFO-Zr | 8.5-9.0 [1, 2] | 33%[3], 85% [2], 88% [1] | 99% | 91 % in serum 37ºC after 24h and 75% in plasma after 24 h# | ~ 1000 per sEV [3] | ~ 6% (well counter)# ~2.5% by PET/CT imaging [1] |
| sEV-L1-Zr | 7.5 | 44% | 100% | Over 95% in plasma after 24 h | ~ 5770 per sEV | ~1.5% (Post mortem) and very low for detection PET-MRI imaging |

### In our experimental settings.
